# Negative linkage disequilibrium between amino acid changing variants reveals interference among deleterious mutations in the human genome

**DOI:** 10.1101/2020.01.15.907097

**Authors:** Jesse A. Garcia, Kirk E. Lohmueller

**Author notes:** Corresponding author: Kirk E. Lohmueller, Department of Ecology and Evolutionary Biology, University of California, Los Angeles, 621 Charles E. Young Drive South Los Angeles, CA 90095-1606.

## Abstract

While there has been extensive work on patterns of linkage disequilibrium (LD) for neutral loci, the extent to which negative selection impacts LD is less clear. Forces like Hill-Robertson interference and negative epistasis are expected to lead to deleterious mutations being found on distinct haplotypes. However, the extent to which these forces depend on the selection and dominance coefficients of deleterious mutations and shape genome-wide patterns of LD in natural populations with complex demographic histories has not been tested. In this study, we first used forward-in-time simulations to generate predictions as to how selection impacts LD. Under models where deleterious mutations have additive effects on fitness, deleterious variants less than 10 kb apart tend to be carried on different haplotypes, generating an excess of negative LD relative to pairs of synonymous SNPs. In contrast, for recessive mutations, there is no consistent ordering of how selection coefficients affect r^2^ decay. We then examined empirical data of modern humans from the 1000 Genomes Project. LD between derived nonsynonymous SNPs is more negative compared to pairs of derived synonymous variants. This result holds when matching SNPs for frequency in the sample (allele count), physical distance, magnitude of background selection, and genetic distance between pairs of variants, suggesting that this result is not due to these potential confounding factors. Lastly, we introduce a new statistic **H_R_**^(j)^ which allows us to detect interference using unphased genotypes. Application of this approach to high-coverage human genome sequences confirms our finding that deleterious alleles tend to be located on different haplotypes more often than are neutral alleles. Our findings suggest that either interference or negative epistasis plays a pervasive role in shaping patterns of LD between deleterious variants in the human genome, and consequently influencing genome-wide patterns of LD.

## Introduction

The non-random association of alleles at different loci is often referred to as linkage disequilibrium (LD). The magnitude of LD between two single nucleotide polymorphisms (SNPs) is shaped by both population processes, such as demographic history, and intrinsic cellular factors, like recombination and gene conversion [1–4]. Previous studies have used LD to estimate demographic parameters of populations such as the historical effective population size (N_e_) and divergence times between populations [2,5,6]. Additionally, methods have been developed to estimate recombination rates from patterns of LD [7–11]. Most of the previous work on modeling patterns of LD has relied on assumptions about selective neutrality among markers. Though some work has been done describing and quantifying the effects of positive selection on patterns of linkage disequilibrium [12–16], the effect of negative selection on genome-wide patterns of LD has received comparably less attention.

Negative selection can influence patterns of LD in two ways: negative synergistic epistasis and Hill-Robertson Interference (HRI). Negative synergistic epistasis [17] occurs when haplotypes carrying multiple deleterious mutations are less fit than predicted by their marginal fitness [17]. This leads to derived deleterious variants being more likely to segregate on distinct haplotypes compared to neutral mutations, creating a deficit of haplotypes containing multiple derived alleles together [18]. Sohail *et al.* showed that rare loss-of-function alleles are underdispersed in human and *Drosophila* genomes [18]. This underdispersion is the consequence of natural selection removing haplotypes containing multiple deleterious alleles, suggesting that nonsynonymous (NS) polymorphisms currently segregating in human and *Drosophila* populations are not only experiencing negative selection, but are also non-independently affecting fitness.

A second way in which negative selection can impact LD is through Hill-Robertson interference (HRI) [19]. In this scenario, one deleterious variant inhibits the removal of a nearby deleterious variant [20, 21]. To understand how this occurs, first, consider a finite population where mutation and genetic drift generate LD between a pair of deleterious variants. Haplotypes carrying multiple deleterious variants (which we will refer to as positive LD) are more effectively removed from this population than haplotypes containing only one deleterious variant because such haplotypes have the lowest fitness. Consequently, selection becomes less efficient at removing haplotypes carrying one deleterious and one non-deleterious alleles (here the deleterious variants are in negative LD) because selection at the first locus conflicts with selection at the second locus. In scenarios with free recombination (r=0.5 per bp), LD between two loci is expected to continue for two generations [20, 21], meaning that selected loci in LD will interfere with each other’s evolutionary dynamics for, on average, two generations. However, when two loci are physically-linked and recombination is low, LD may persist longer allowing the linked loci to interfere with each other’s evolutionary dynamics for longer periods of time, thus reducing the effectiveness of selection at both sites. HRI predicts pairs of deleterious SNPs will exhibit negative LD, especially when they are separated by small genetic distances, i.e., with recombination frequencies on the order of one centimorgan (cM) [22–24].

Hill and Robertson [19] first studied the effect of linkage on limits to natural selection with only two additive loci under natural selection. They reported that interference creates a detectable excess of negative LD with biologically relevant effective population sizes and variants with realistic selection coefficients. However, their simulations and theory did not incorporate new mutations or multiple loci that differed in age and distance from focal mutations. Adding to their work, McVean and Charlesworth [22] studied the effects of Hill-Robertson interference between weakly selected mutations. They simulated multiple weakly selected mutations and observed that weak selection Hill-Robertson interference (wsHR) generates negative LD amongst beneficial mutations. This excess of negative LD was observed to be most apparent among alleles that were physically close to each other and appeared to have a negative relationship with distance. In their paper, they suggest that interference is a prevalent force driving distribution of biased codon usage in *Drosophila*. Although they looked at the effects of interference caused by negative selection on fixation probabilities, heterozygosity, and average time to loss, they did not examine the impact of interference amongst deleterious sites on LD. Additionally, Comeron and Kreitman showed via simulations that interference among multiple positively selected variants should create an excess of negative LD among low frequency variants while reducing overall levels of neutral polymorphism [23]. However, these simulation studies considered only populations of constant size, and their applicability to the LD patterns of natural populations with more complex demographic histories and multiple sites under negative selection remains understudied. Indeed, recent work has found that the expected pattern of background selection is heavily affected by the demographic history of the population [25, 26].

Despite there being decades of theoretical work on Hill-Robertson interference, there has been comparatively less work quantifying it in natural populations. Comeron *et al.* hypothesized that interference could influence the spatial distribution of putatively beneficial codons [27]. Using forward simulations and biologically relevant recombination rates, Comeron *et al.* proposed that a multi-site model of a finite population, with mutations, selection, and linkage could predict the observed relationship between magnitude of codon usage bias and coding sequence length observed in *Drosophila*.

The role of interference has also been assessed in primates (human, chimpanzee, and rhesus macaque) by quantifying the relationship between recombination and d_N_/d_S_ [28]. Bullaughey *et al.* (2008) suggested that there is no detectable effect of recombination on rates of protein evolution. Later, Hussin *et al.* [29] looked for signatures of interference by quantifying the relative enrichment of deleterious mutations in cold spots of recombination. Theory suggests that cold spots of recombination should be enriched with slightly deleterious mutations relative to hotspots of recombination [20, 30]. They identified this relative enrichment in exons, though the strength varied across populations. Additionally, they observed that conserved exons in recombination cold spots are enriched with haplotypes with two NS variants relative to exonic haplotypes in hotspots of recombination. Hussin *et al.* concluded this excess of mutational burden in cold spots would be expected under a process similar to Muller’s ratchet [30]. Here, the least-loaded haplotype class cannot be quickly regenerated if recombination is scarce because it is degraded by drift or new mutations. These results suggested that interference might play a significant role in determining patterns of genetic diversity in human autosomes. Although this study examined the effects of negative selection on the distribution of burden and enrichment of deleterious variation across haplotypes and recombination rates in the human genome, it did not explicitly study the effect of negative selection on LD summary statistics (*r*^2^, *D*, *D*’).

In the present study, we first examined how negative selection affects levels of LD among deleterious NS SNPs relative to the LD among neutral synonymous (S) SNPs using forward simulations. Although all summary statistics of LD are dependent on allele frequency, we control for this possible confounder in our comparisons by limiting pairwise LD calculations using the method of frequency matching described by Eberle et al. [31]. We find in forward simulations with a human-oriented recombination rate, mutation rate, and distribution of fitness effects of new mutations, that negative selection induces a detectable excess of negative LD among derived SNPs. Next, we used human data from the Phase 3 1000 Genomes Project (1KGP) [32] and tested for a difference in LD patterns between NS and S SNPs. We found that pairs of derived NS variants have more negative LD between them compared to matched pairs of derived S variants. Additionally, to replicate our results, and to provide a method to detect interference in unphased data sets, we introduce a new summary statistic H_R_^(j)^. Using this statistic, we demonstrate signatures of interference in the New York Genome Center’s (NYGC) unphased high coverage resequencing of 1KGP individuals [33]. Our findings of an excess of negative LD between pairs of putatively deleterious variants suggest that either interference or negative synergistic epistasis play a pervasive role in shaping patterns of LD between proximal NS variants in the human genome.

## Results

### Forward Simulations

We performed forward simulations using SLiM 3.0 [34]. Each generated chromosome was approximately 5 Mb long and contained intergenic, intronic, and exonic regions. Only NS mutations within exonic regions experienced negative selection (see Methods). We simulated under three demographic models. Model 1 consists of a population of 10,000 individuals evolving for 100,000 generations. Model 2 is the model of human demography by Gravel *et al.* [35] and implemented into SLiM by Haller and Messer [34]. Model 3 is identical to Model 2 except that there is no migration across the populations (**SI Fig 1**). For each simulation replicate, we sampled 50 individuals from the African population and computed the various LD statistics as described in Methods.

### Predicted effect of negative selection on linkage disequilibrium decay

To gain an overall understanding of how negative selection can impact LD patterns, as measured by the squared correlation coefficient between SNPs (*r*^2^), we first simulated populations under demographic Model 1 (constant-size population) where NS mutations all have the same effects on fitness (Fig 1). The strength of negative selection, degree of dominance, and recombination rate, all interact and affect the mean r^2^ decay with physical distance between SNPs (Fig 1). This effect is most visible in simulation replicates with low recombination rates. Specifically, under the additive model, i.e., where *h* = 0.5, simulations with weakly to moderately deleterious (−0.001<*s*<-0.01) mutations have lower values of *r*^2^ than do simulations with only neutral variants. However, simulations with deleterious variants with *s*= −0.1 tended to have the highest mean *r*^2^ for a given physical distance, mirroring the LD patterns of neutral variants. This is likely in part due to the fact that strongly deleterious mutations are not expected to segregate within a population for long. Therefore, the effects of interference are diminished [24].

**Fig 1.**
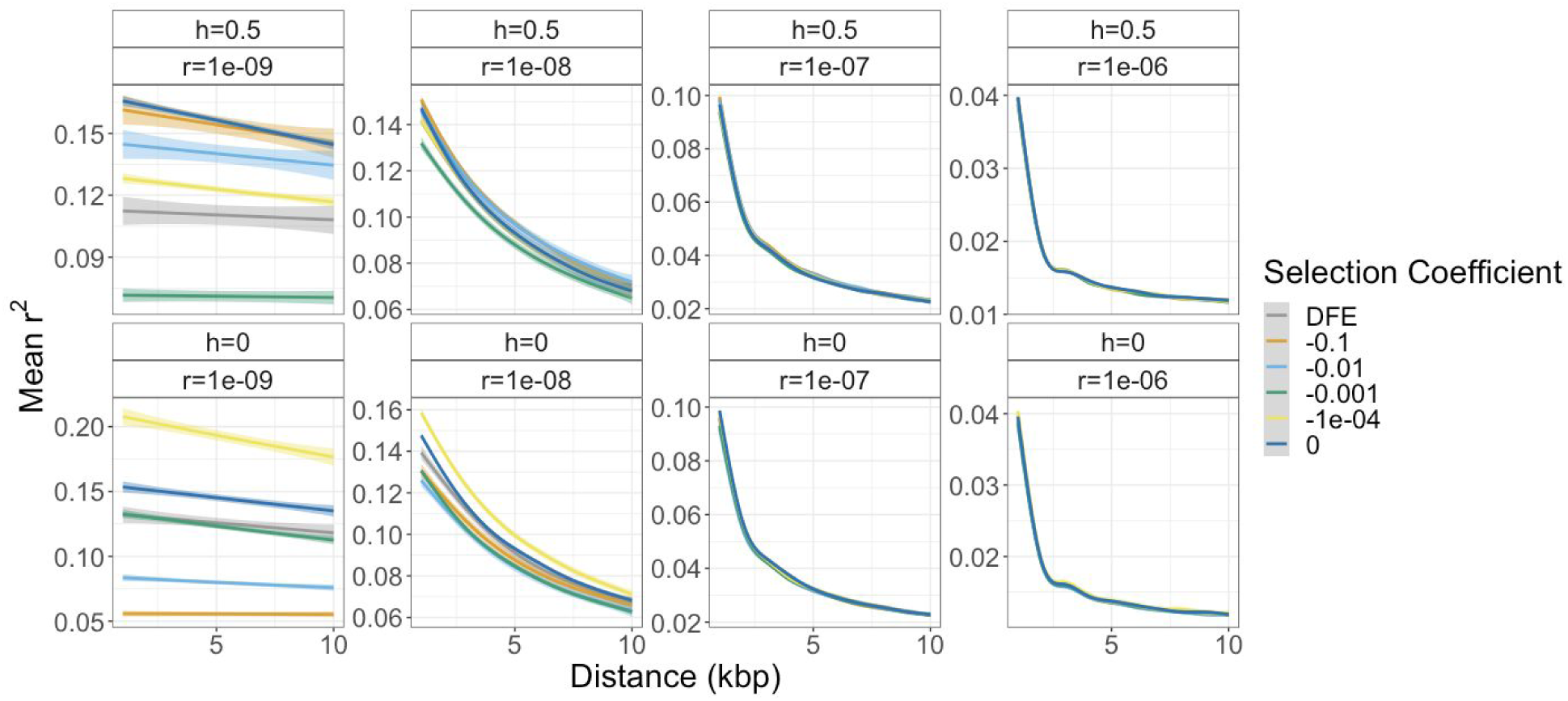
Decay of mean *r*^2^ for different recombination and dominance parameters. The differences in the LD decay curves are most apparent with recombination rate *r*=1 x 10^-9^ per bp and depend on the dominance coefficient (*h*) of mutations and the selection coefficient of NS mutations. DFE curves come from simulated populations where new NS mutations have selection coefficients following our defined distribution of fitness effects (see text).

For simulations with recessive deleterious mutations (*h*=0), there is a complex relationship between the selection coefficient and LD decay (Fig 1). Here moderately and strongly deleterious (*s*=-0.01 and −0.1) variants have the lowest values of *r*^2^ at a given physical distance. However, simulations with NS variants with selection coefficients of −0.0001 have the largest mean r^2^. Thus, for the recessive case, there is no consistent ordering of how selection coefficients affect genome-wide *r*^2^ decay with physical distance.

For simulated genomes with recombination rates greater than 1 x 10^-9^ per bp, the effect of negative selection on mean *r*^2^ and its decay is less pronounced. Here, simulations with deleterious variants that arise with drastically different selection coefficients displayed similar *r*^2^ decay patterns. Furthermore, as the recombination rate increased, the overall magnitude of *r*^2^ decreased, as expected given previous work [19, 36].

### LD of doubletons under direct negative selection

The simulations described above analyze the overall patterns of LD among all pairs of variants within each simulation replicate. As such, they included both neutral (S) and negatively selected (NS) variants. Additionally, those *r*^2^ computations included all variants, regardless of allele frequency. It is well-known that LD summary statistics are influenced by allele frequency [3, 31]. Therefore, we controlled for this effect in the subsequent sets of simulations by only considering pairs of SNPs with the same frequency in our samples. Further, we only calculated LD statistics between pairs of SNPs with the same functional annotation. For example, we computed LD only between pairs of NS SNPs or between pairs of S SNPs.

We begin by focusing on doubletons (derived variants that show up in our sample of 50 diploid individuals twice, or variants with a frequency in our sample of 2/100) because we hypothesized that doubletons would be enriched with polymorphisms that are more strongly influenced by negative selection relative to higher frequency variants [37–39]. Consistent with this hypothesis, simulations (see Methods) show that deleterious NS variants with a frequency greater than 2/100 in a sample of 50 individuals tend to be less deleterious than doubletons (**SI Fig 2**). Additionally, relative to singletons, we expect doubletons to be less influenced by sequencing error or errors in variant calling in empirical data. We quantify LD between derived doubletons using *D* rather than *r*^2^ because the sign of *D* is informative regarding whether the derived alleles preferentially occur on the same haplotypes (are in coupling, also called positive LD) or different haplotypes (are in repulsion, also called negative LD) (see **SI Fig 3**). If a pair of NS doubletons both occur on the same haplotype in our sample of 100 chromosomes, they will have a *D* value of 0.0196. Pairs of derived NS doubletons that never occur on the same haplotype in our sample have a *D* value of −0.0004. Thus, the average value of *D* at a given distance reflects the number of pairs of SNPs that occur on the same haplotype (more positive) or on different haplotypes (more negative).

Using the same simulations as described above, we found that *D* between doubletons is greatly affected by negative selection (Fig 2). When assuming additive effects on fitness, *D* for weakly and moderately deleterious (*s*=-1e-04, *s*=-1e-03, *s*=-1e-02) variants is more negative than those for neutral doubletons. Thus, while the LD patterns depend on the specific selection coefficients, dominance coefficients, and recombination rates, in general, deleterious NS doubletons (s<0) tend to occur on different haplotypes more often than S SNPs that are neutral in these simulations (dark blue curve in Fig 2). Similar trends are seen when computing other summary statistics of LD (e.g. *D*’ and *r*^2^, see **SI Fig 4**).

**Fig 2.**
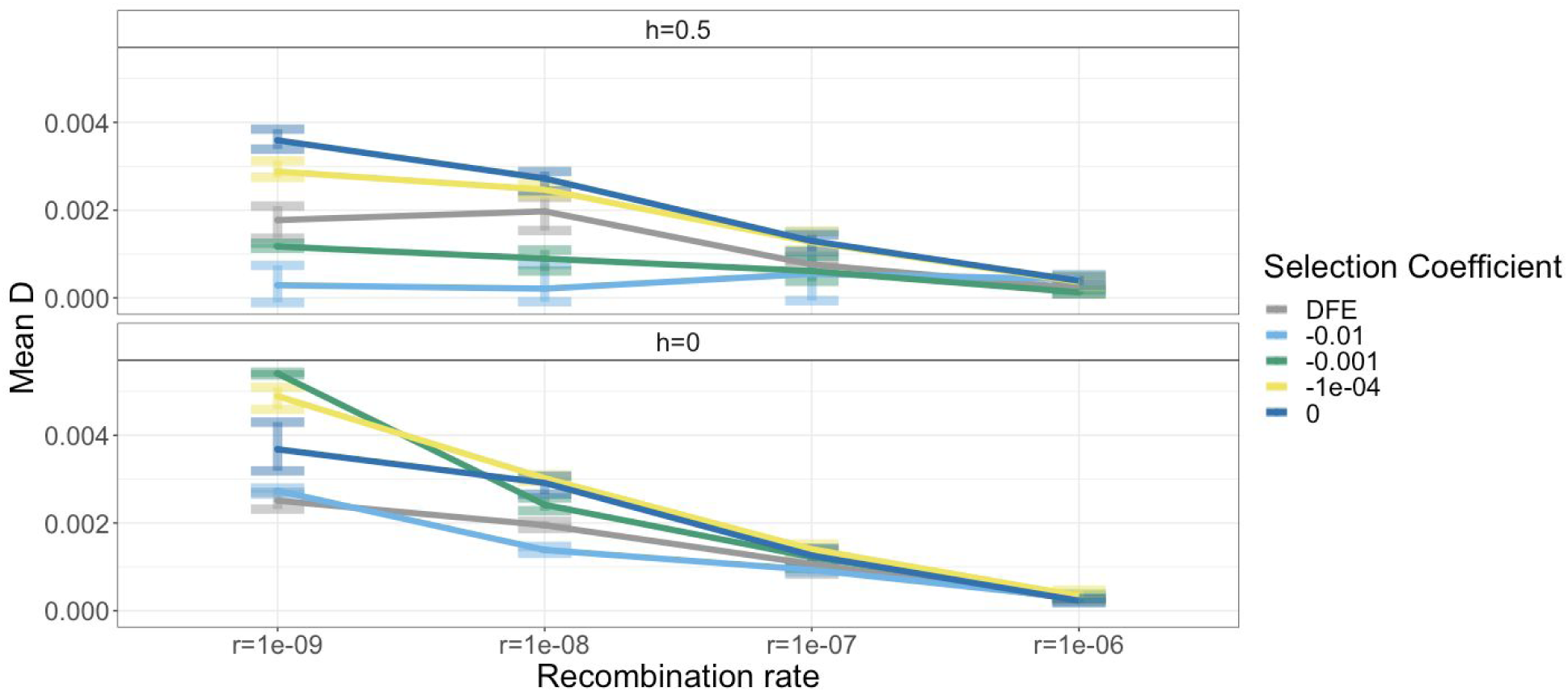
Mean *D* for simulated NS doubletons within 10kb from each other across different recombination and dominance parameters. The differences in the decay curves are most apparent for recombination rate *r*=1 x 10^-9^ per bp and depend on the dominance coefficient of mutations (*h*) and the selection coefficient of NS mutations.

For recessive variants, pairs of moderately deleterious (*s*=-0.01) doubletons also tend to have more negative values of *D* than pairs of neutral doubletons. However, more weakly deleterious SNPs (*s*=1e-04 and *s*=1e-03) have mean values of *D* that are greater than those values for neutral SNPs (Fig 2). These patterns appeared to hold across all recombination rates, though the magnitude of the difference is greatest in regions with low recombination rates.

A main focus of this study is to describe the effects of negative selection on patterns of LD in the human genome. Thus, we next simulated human-like exomes with selection coefficients of NS mutations coming from a distribution of fitness effects inferred from human population genomic data (see DFE in Methods). These simulations suggest that human-like levels of negative selection are predicted to generate negative LD amongst deleterious variants relative to the LD observed of neutral mutations with the same demography and recombination rate (gray curve in Fig 2).

In the previous section, we show that negative selection is expected to influence LD patterns compared to a scenario where all mutations are neutral. However, in real genomes, putatively neutral S and intronic variants are interspersed with deleterious NS mutations. We next compared LD patterns between pairs of NS SNPs (with *s*= −0.0001) to the LD patterns of pairs of neutral S SNPs (no effect on fitness) in our simulations. Consistent with our previous simulations, pairs of derived NS doubleton SNPs (blue line) have more negative LD than pairs of derived S doubletons (yellow line) (**SI Fig 5**). This finding predicts that NS doubletons tend to be located on different haplotypes more often than S doubletons.

As the degree of LD between a pair of doubletons also is influenced by the amount of recombination that occurs between them, it is possible that there could be different genetic distances (cM) between pairs of NS and S SNPs. To test for this, we simulated 50 replicates under neutrality but with the same distribution of exons and introns. We simulated NS SNPs ‘without negative selection’ by simulating mutations with a NS mutation rate, but with *s*=0. In summary, the LD patterns for the pairs of NS SNPs without negative selection (red line) matches the LD patterns of pairs of S SNPs (yellow and purple line) (**SI Fig 5**). These results suggest that the distribution of functional elements is not responsible for the differences in LD patterns between deleterious NS SNPs and interspersed neutral SNPs.

We next examined how negative selection and interference are expected to affect LD patterns around pairs of putatively neutral markers interdigitated with and potentially linked to negatively selected mutations. We compared the LD decay curve of neutral S variants in simulations with deleterious NS mutations, to the LD decay curve of S variants in simulations where no deleterious variation was present. When pairs of SNPs are close together (< 500bp apart), the mean *r*^2^ for S SNPs in simulations with negative selection was slightly higher than that for S SNPs in simulations without negative selection, i.e., completely neutral evolution (**SI Fig 5**). These results suggest that not only does negative selection affect the LD of negatively selected variants themselves, but it also impacts the magnitude of LD between linked neutral variants.

### Effect of negative selection and complex demography on LD

We next examined how complex, non-equilibrium demography impacts LD between deleterious variants. We hypothesized that although population growth, and migration all perturb levels of LD, we should still be able to detect interference by comparing LD between pairs of deleterious NS variants to the LD between pairs of neutral S variants. To test this, we simulated under Model 2, which is a more realistic model for human demography, containing exponential population growth, and migration (**SI Fig 1**). These simulations also use a distribution of fitness effects for deleterious mutations (see Methods). Here we still see the effects of interference generating an excess of negative LD amongst derived deleterious doubletons as NS doubletons have lower values of D than do neutral S doubletons (**SI Fig 6**). Again, the difference between functional annotations (NS or S) is greater for SNPs at low recombination rates. These findings suggest that the relative excess of negative LD can be detected in human-like demography and is not limited to populations with a constant long-term N_e_ and in equilibrium.

We hypothesized that migration across populations might disrupt the strength of the correlation between allele frequency and fitness effects, which in turn could affect the strength of interference and LD. In Model 2, low frequency variants in a sample might be at low frequency due to having just recently arrived in the population via migration. These low-frequency variants could be at higher frequencies in other populations giving them a high probability of migrating to a new population. As such, these doubletons may be at low frequency due to the fact that they have recently arrived to a new population, rather than being actively kept at low frequency by negative selection. To test for this effect, we looked at the relationship between the mean selection coefficient of doubletons based on their population of origin (**SI Fig 7**). Although we sample from the African population, there are variants in the sample that arose in other populations. On average, variants that originated from different populations then migrated into Africa are less deleterious (**SI Fig 7**). Additionally, we hypothesized that, if migration is the source of less striking differences in LD between NS and S variants in Model 2, then the same demographic model without migration across populations will show a larger difference in LD patterns between neutral and deleterious SNPs. To test this hypothesis, we conducted additional simulations under the same demographic history as Model 2, though without migration (Model 3). Model 3 shows a slightly greater excess of negative LD among derived NS variants compared to S variants (**SI Fig 6** dashed lines) than that seen under Model 2 (**SI Fig 6** solid lines). However, because this pattern is quite subtle, it suggests that migration has had a limited impact on differences in patterns of LD across different types of SNPs.

Additionally, in our simulations, we quantified differences in the proportion of pairs of NS and S doubletons in complete repulsion (i.e. D’ = - 1; **SI Fig 8)**. We hypothesized that the more negative LD statistics of simulated deleterious NS variants is due to more pairs of NS variants being in complete repulsion (D’ = - 1) than were pairs of S variants. In our study, pairs of doubletons in complete repulsion refers to the case where a pair of doubletons are distributed across haplotypes in a way such that derived variants never co-occur on the same haplotype in our sample.

For example, in the complete repulsion case for a pair of biallelic doubletons annotated as NS, there are two segregating loci. We denote the first locus as either being “a” (ancestral allele) or “A” (derived allele). Similarly, the second locus can be denoted with either “b” (ancestral allele) or “B” (derived allele). Haplotypes that contain ancestral variants at both loci would occur in 96 of the 100 haplotypes in our sample (n_ab_=96), haplotypes that contain one ancestral variant and one derived variant could occur four times out of 100 (n_Ab_ + n_aB_ = 4), and haplotypes that contain derived variants at both loci could occur zero times (n_AB_=0). In the simulations in which we quantify interference, we hypothesized that negative selection will act to remove AB haplotypes of NS (deleterious) doubletons and not remove AB haplotypes of S (neutral) doubletons. Additionally, negative selection will more rapidly remove AB haplotypes of pairs of NS variants relative to Ab or aB haplotypes. We tested this prediction in our forward simulation framework and found that for low recombination rates, more pairs of NS doubletons were in complete repulsion than were pairs of S doubletons. This suggests that haplotypes carrying two deleterious alleles are most efficiently selected out from the population, and those carrying only one deleterious allele end up persisting in the population. These results are predicted under Hill-Robertson interference [19, 21].

### A new statistic to quantify the effect of selection on the distribution of unphased 2-locus genotypes (H_R_^(j)^)

Thus far we have examined traditional LD statistics that require phased haplotype information. We typically do not have this information and instead have to rely on computational phasing. Here we consider the case where phasing is not available, and present a statistic quantifying whether derived alleles are more likely to be coupled (i.e. on the same haplotype) or in repulsion (i.e. on different haplotypes) for different functional annotations.

Consider a pair of singleton variants. An individual can be homozygous ancestral (0/0,0/0) at both variants, heterozygous at one SNP (0/1,0/0 or 0/0,0/1), or doubly heterozygous (0/1,0/1). If an individual is heterozygous at only one of the SNPs, then by definition, the two singletons must be on different haplotypes. If an individual is heterozygous for both SNPs, it is possible that both derived alleles are carried on the same haplotype. Alternatively, if an individual is heterozygous for both SNPs, it is possible that both derived alleles are carried on different haplotypes (see **SI Fig 3**). Quantifying the distribution of homozygous ancestral, singly heterozygous, and doubly heterozygous genotypes does not require information about haplotype phase and thus can be applied to genotype data. We formalize this idea in a statistic that we call **H_R_**^(j)^ (see Methods for a more detailed description and how to compute **H_R_**^(j)^; also see **SI Fig 9**). Our statistic can be calculated from pairs of derived variants at different frequencies in a sample. Applying this statistic to only doubletons in a sample of 50 diploid individuals, corresponding to variants with frequency of 2/100, we call it H_R_^(2)^. Essentially, **H_R_**^(j)^ counts the average number of individuals heterozygous at both SNPs for pairs of SNPs at a given distance. Doubletons in a sample of 50 are not often found in a homozygous state. In order for this to happen, a variant must appear twice in a sample, but in only one individual. If an individual is homozygous derived at both SNPs, they are not counted in **H_R_**^(j)^. Based on predictions from our simulations using other LD statistics (*D*, *D*’, *r*^2^), we hypothesized **H_R_**^(j)^ to be lower for deleterious SNPs than for neutral SNPs. To test this hypothesis and examine the behavior of the **H_R_**^(j)^ statistic, we simulated under Model 2 across two different recombination rates (r=1 x 10^-9^ per bp and r=1 x 10^-8^ per bp). Our simulations show that unphased genotypes and H_R_^(2)^ are affected by negative selection. Specifically, for both recombination rates (Fig 3) there is a depletion in mean H_R_^(2)^ and H_R_^(1)^ for pairs of NS variants compared to S variants. Thus, we conclude that H_R_^(j)^ can detect interference between deleterious variants using unphased genotypes.

**Fig 3.**
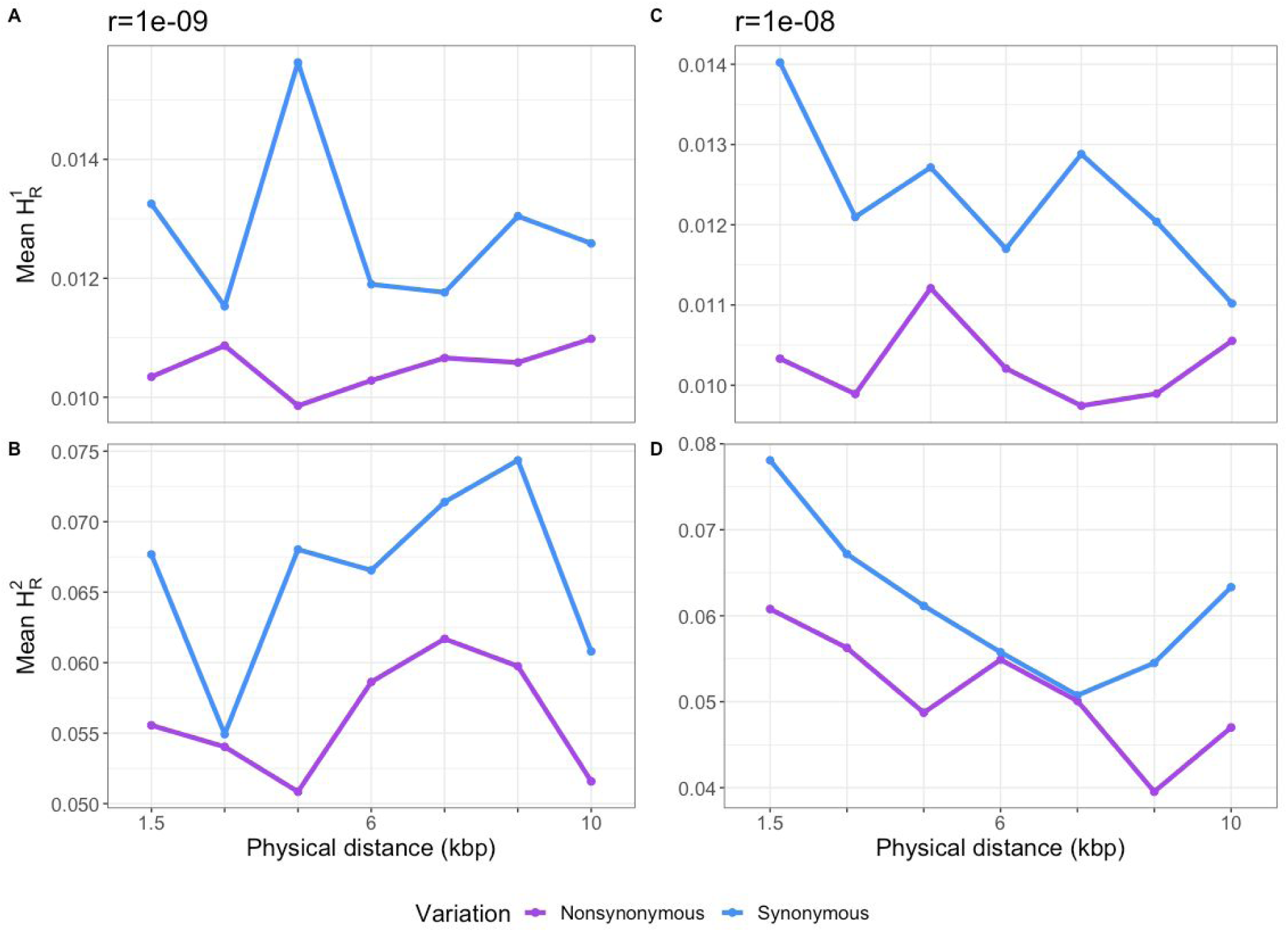
Distribution of mean H_R_^(1)^ and H_R_^(2)^ across simulated data with different recombination rates and selection coefficients of NS mutations drawn from a DFE. Forward simulations predict NS pairs of variants have a mean H_R_^(j)^ that is on average lower than that of S pairs of variants. (A) H_R_^(1)^ for simulations with a low recombination rate (*r*=1 x 10^-9^ per bp). (B) H_R_^(2)^ for simulations with a low recombination rate. (C) H_R_^(1)^ for simulations with average recombination rate (*r*=1 x 10^-8^ per bp) (D) H_R_^(2)^ for simulations with an average recombination rate. Each point is the mean H_R_^(j)^ statistic in a 1.5kbp wide distance bin.

### Comparison of LD between NS and S SNPs in humans

We next tested whether negative selection has affected patterns of LD across the human genome. Here we examined 50 diploid genomes from the Yoruban population of the 1KGP [32]. We first computed *r*^2^, *D*, and D’ for pairs of low-frequency (in this study frequency equal to or less than 5/100, or allele count equal to or less than five) NS variants matched for allele frequency. The same was done with pairs of low frequency S variants. To test for a significant difference in the amount of LD between S and NS SNPs, we developed a permutation test where we matched each pair of NS SNPs to a pair of S SNPs that had the same derived allele frequency, physical distance between the pair of SNPs, genetic distance between the pair of SNPs, and magnitude of background selection (See Methods, **SI Fig 11**).

We observed a qualitative difference in the mean *D*’ of matched pairs of NS and S variants across genetic distance bins (Fig 4a). Specifically, NS SNPs have less LD at a given distance, as reflected by the lower values of *D*’, compared to S SNPs. Application of the matched-pairs permutation test showed that the observed difference of mean *D*’ between NS and S variants is significant (p < .0005) (Fig 4b). Likewise, other LD summary statistics show lower mean values for pairs of NS SNPs than S SNPs (Fig 4b). To evaluate the performance of our matched-pairs permutation test, we simulated genomes under a scaled version of Model 1 with simulations that: 1) included negative selection using the distribution of fitness effects inferred by [28], and 2) did not contain negative selection. While a significantly more negative mean *D*’ was seen in simulations with negative selection (Fig 4c left histogram), this significant difference was not seen in simulations without negative selection (Fig 4c right histogram). Additionally, in the simulations without negative selection, we performed this matched-pairs permutation test on other summary statistics of LD and observed no significant difference between pairs of NS variants and S variants (**SI Fig 12)**. Because of our matched-pairs permutation procedure, we conclude this excess of negative LD between pairs of NS SNPs was not due to differences in local recombination rates or differences in background selection.

**Fig 4.**
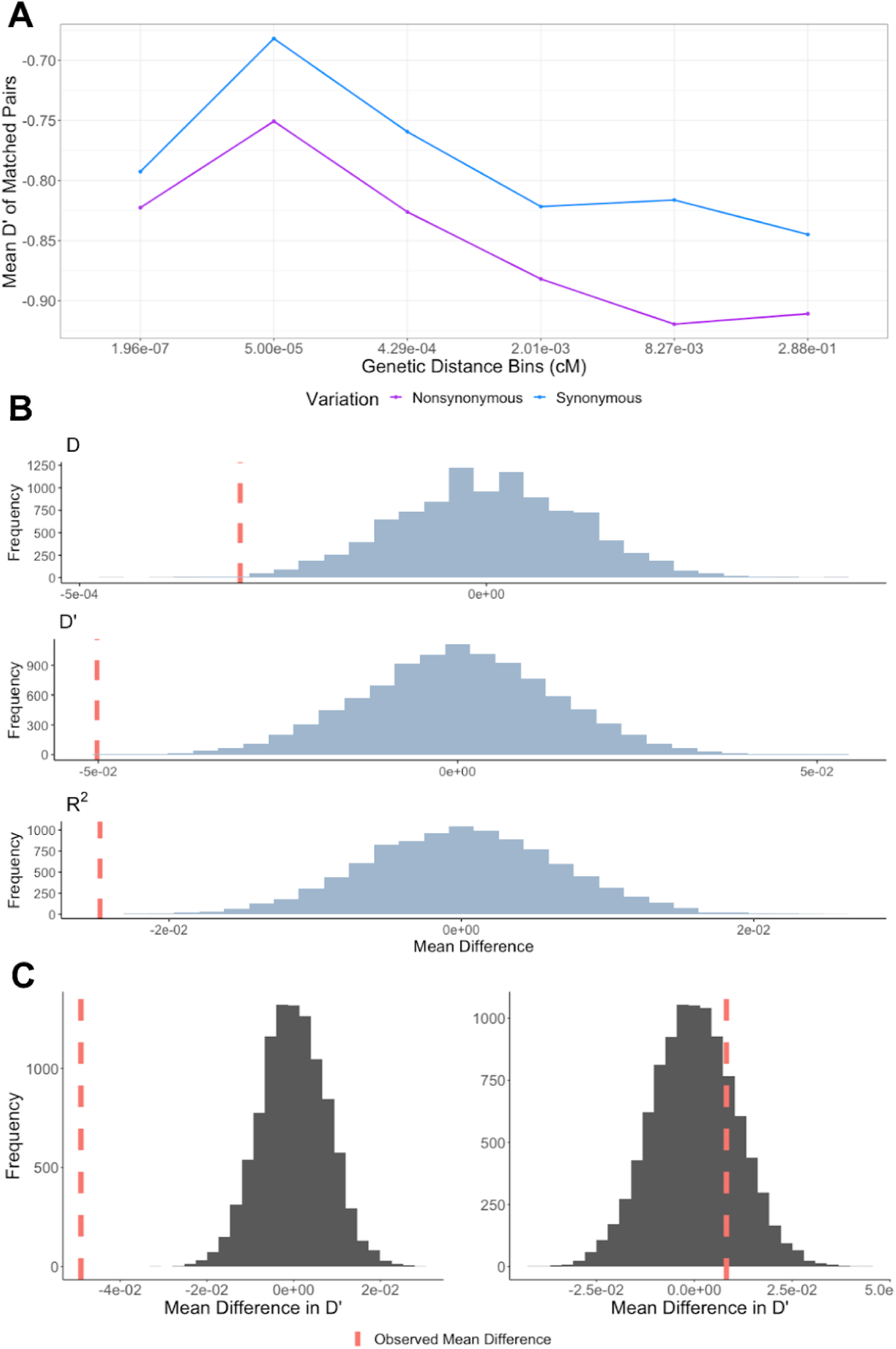
Pairs of NS variants have lower LD compared to pairs of S variants in human genetic data. (A) Pairs of low frequency (variants with minor allele count <=5 in a sample of 50 individuals) NS variants in the YRI population of the 1KGP have a more negative mean *D*’ compared to pairs of S variants across genetic distance bins. (B) In the YRI population of the 1KGP, NS pairs of variants have a significantly lower *D*, *D*’, and *r*^2^ compared to matched pairs of S variants. (C) In simulations with a biologically relevant DFE to humans (left histogram), a matched-pairs permutation test can detect difference in *D* between pairs of NS and S variants. In simulations with no negative selection (right histogram), a matched-pairs permutation test does not detect a difference in *D* between pairs of NS variants and pairs of S variants. This suggests negative selection can explain the reduction in LD we see in the NS SNPs in the 1KGP data.

### Analysis of additional populations

We also tested for differences in LD patterns between pairs of NS and S variants in other human populations. We selected data from 50 individuals from four populations included in the 1KGP (Han Chinese in Beijing, China (CHB), Utah Residents with Northern and Western European Ancestry (CEU), Mexican Ancestry from Los Angeles USA (MXL), Japanese in Tokyo, Japan (JPT)) and performed identical analysis to those described above to quantify the difference in LD between the different types of SNPs. As in the YRI population, we observed that mean *D*’ between matched pairs of NS SNPs was clearly lower than that for pairs of S SNPs across genetic distance bins in three out of four populations (Fig 5a). Furthermore, we found a significant excess of negative LD amongst matched pairs of NS variants relative to S variants in three out of four populations (CHB, MXL, JPT) using our matched-pairs permutation test (Fig 5b). In the CEU, there was still more negative *D*’ for pairs of NS variants (-.7806) compared to pairs of S variants (−0.7722), though the difference was not significant. These findings illustrate that across several human populations, NS derived alleles tend to co-occur on different haplotypes more frequently than do S variants that have the same frequency.

**Fig 5.**
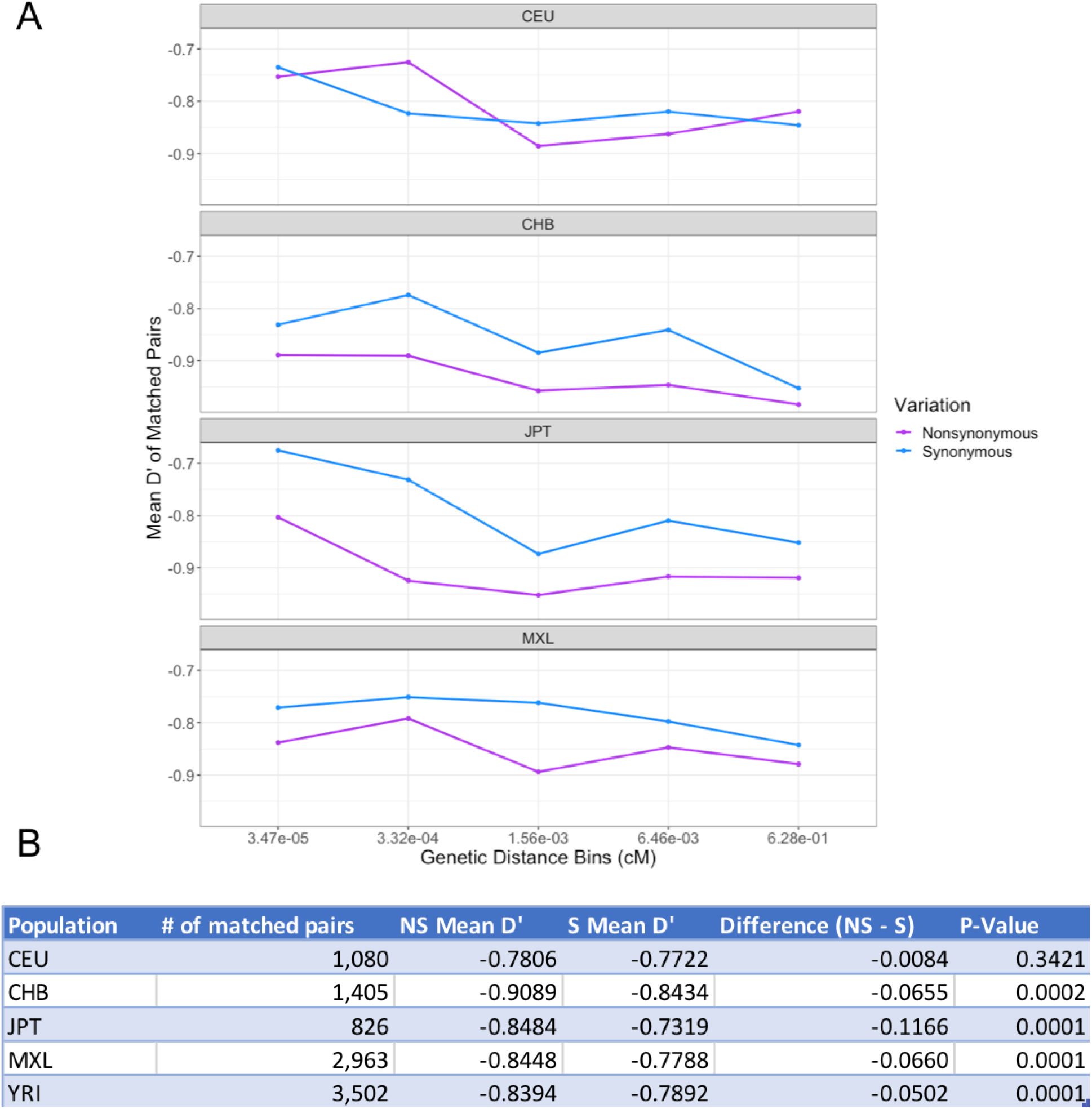
Results from the matched-pairs permutation test for 1KGP populations. (A) Across four other tested population in the 1KGP, NS pairs of variants have lower values of *D*’ than their matched S counterparts. (B) Matched-pairs permutation test shows significantly more negative average LD between pairs of NS SNPs than S SNPs in all populations except the CEU population, where this difference is not significant.

### Replication in high-coverage sequence data

Our previous analysis used 1KGP Phase 3 data which is the product of genotype imputation and statistical phasing. To mitigate the possible effects of imputation and haplotype phasing errors on our analysis, we also examined the distribution of NS and S variants using unphased genotypes. In addition to using the 1KGP Phase 3 dataset described previously, we also used the 30X WGS of the 1000 Genome Project samples, sequenced by the New York Genome Center and funded by NHGRI (http://ftp.1000genomes.ebi.ac.uk/vol1/ftp/data_collections/1000G_2504_high_coverage/working/20190425_NYGC_GATK/) [33].

Our simulations show that H_R_^(2)^ and H_R_^(1)^ should be lower for pairs of NS variants compared to paris of S variants (Fig 6). In the 1KGP Phase 3 YRI data, on average, pairs of NS variants have a lower mean H_R_^(2)^ and H_R_^(1)^ (Fig 6ab) compared to S variants. This difference is statistically significant for both doubletons (observed difference = −0.017, permutation test p = 0.011) and singletons (observed difference −0.006, permutation test p = 0.001). Importantly, the 30X WGS of the same 1000 Genome samples sequenced to higher coverage by the New York Genome Center replicates this result (Fig 6cd). In this data set, the observed difference in mean H_R_^(2)^ for NS and S variants is −0.065 (p = 0.0088), and the observed difference in mean H_R_^(1)^ is −0.018 (p = 0.001). Thus, our finding that NS SNPs tend to be located on different haplotypes more often than S SNPs is robust to the specific dataset used and is not due to phasing and imputation errors in the 1KGP data.

**Fig 6.**
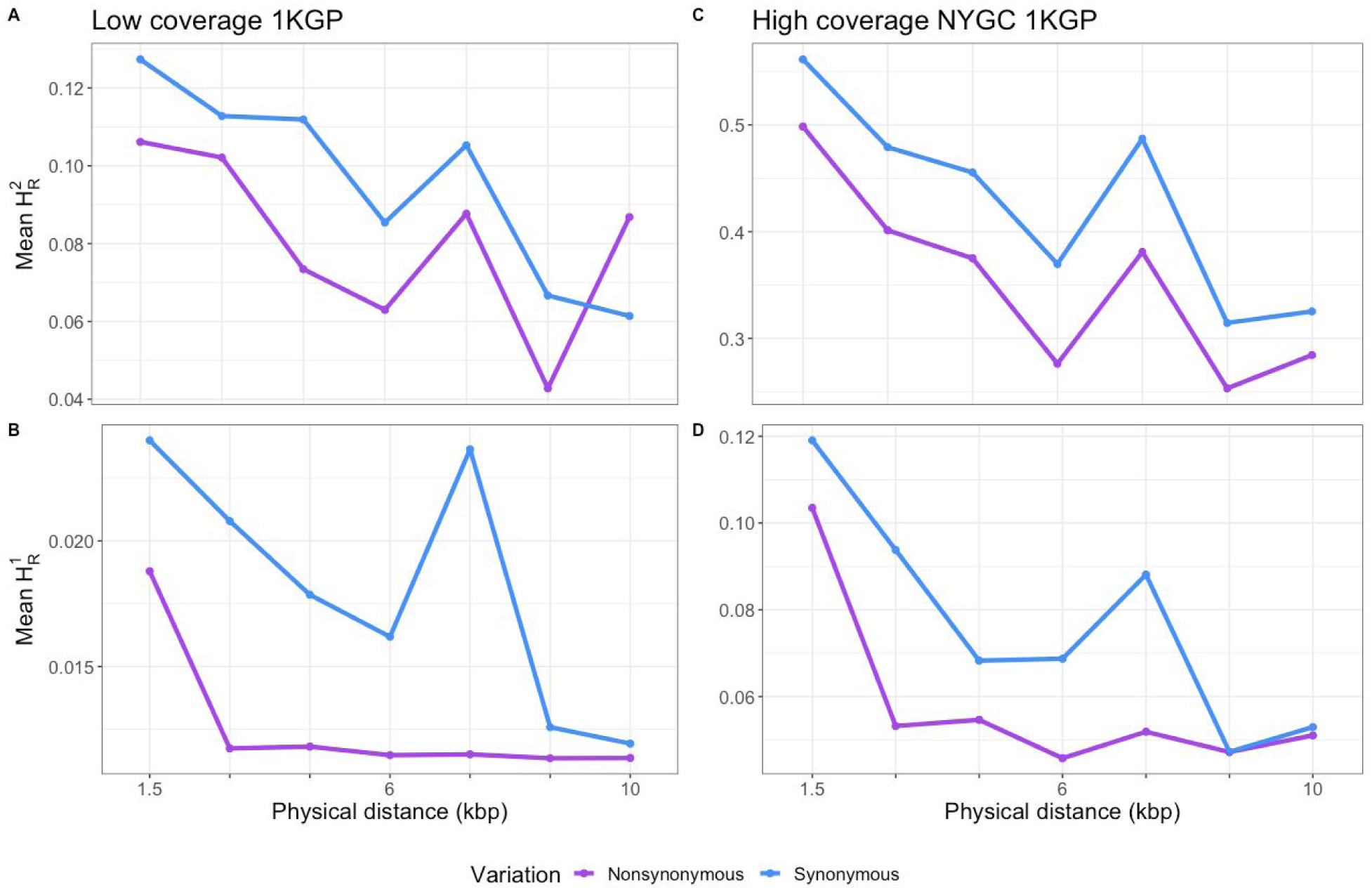
Distribution of mean H_R_^(2)^ and H_R_^(1)^ across empirical data sets with different coverages. (A, B) Low-coverage 1KGP data. (C,D) High-coverage NYGC 1KGP data. For both datasets, mean H_R_^(2)^ and mean H_R_^(1)^ between NS pairs of doubletons is on average lower than that of S pairs of variants.

### Can Hill-Robertson effects account for more negative LD between NS SNPs?

We hypothesize that Hill-Robertson effects are more likely to impact pairs of variants close to each other, whereas other forces, like synergistic epistasis are thought to act on pairs of variants further apart [18]. To better compare the difference between NS and S SNPs as a function of distance, we normalized the difference in *D* between NS and S pairs of SNPs by dividing by the mean *D* of S SNPs (Fig 7). Overall, this statistic is negative in the empirical data across the entire range of genetic distances considered (green line in Fig 7). Simulations under Model 2 show a similar pattern, except that the normalized *D* tends toward 0 with increasing genetic distance between SNPs. To assess whether the difference between the simulations and empirical data could be due to the fact that the empirical data has fewer pairs of SNPs (14,608) than do the simulations, we resampled 14,608 pairs from the simulations with the same frequency as those in the empirical data. Here, the normalized difference in LD calculated from the empirical data falls within the range of values seen in the simulations. Thus, simulations including negative selection can recapitulate the difference in LD between NS and S SNPs at different intervals of genetic distance. We also examined the relationship of mean normalized difference in *D* in CEU and CHB and observed a similar relationship with genetic distance (**SI Fig 13**).

**Fig 7.**
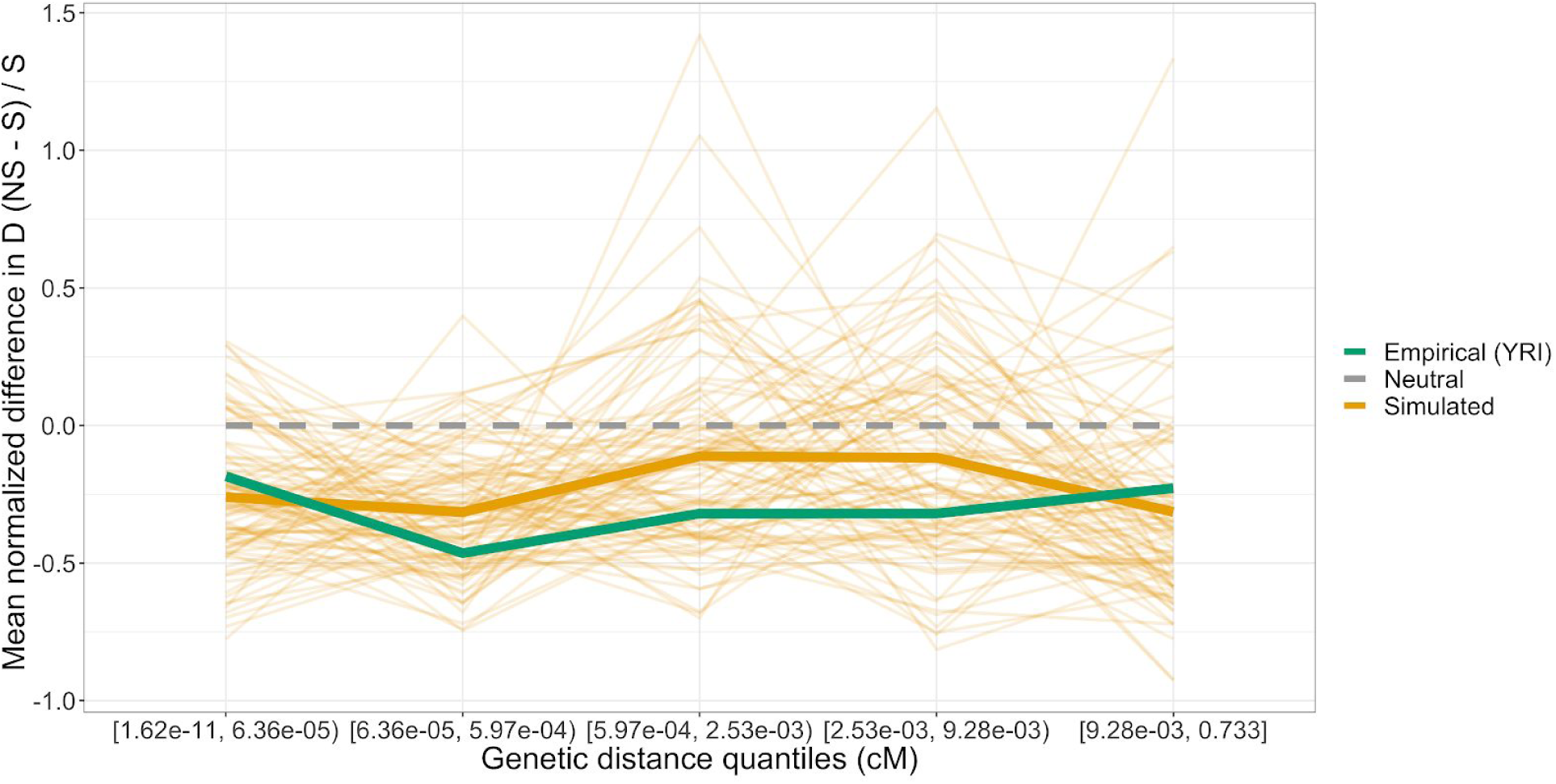
Mean normalized difference in *D* across genetic distance quantiles. Empirical (green) and simulated data shows a deficit in *D* between NS variants. The lighter orange lines show 100 resamples of the simulated data. Each resample of the simulated data has the same amount of variants with the same allele count as the empirical data.

Since our forward simulations consisted of mutations that only affect fitness multiplicatively across loci, we conclude that the excess of negative LD between NS SNPs can be explained by interference. However, we cannot rule out a contribution from synergistic epistasis in the empirical data (see Discussion).

## Discussion

Here we have shown that the dynamics of LD are influenced by the magnitude and dominance of fitness effects of deleterious mutations, the recombination rate, and the underlying demographic history of a population. In our simulations we show how Hill-Robertson interference is expected to influence LD patterns between deleterious mutations and find that the effects are likely to be stronger in regions of lower recombination (Fig 1 and Fig 2), consistent with previous work [19,22,29]. Using a human demographic model and DFE, our simulations suggest that interference should be detectable in both low recombination regions (*r*=1 x 10^-9^ per bp) and regions with an intermediate recombination rate (*r*=1 x 10^-8^ per bp) in the human genome (**SI Fig 6** and Fig 3). We then searched for these patterns in several genomic datasets from several human populations and with different types of data (low-coverage and exome sequence data followed by imputation as well as high-coverage whole genome sequence data). Overall, for most of the comparisons considered, NS SNPs appear to be present on different haplotypes more frequently than S SNPs having the same allele frequency (Fig 4 and Fig 5).

To quantify the excess of negative LD among NS variants relative to S variants, we implemented a matched-pairs permutation test. When we applied this method to human data from the 1KGP, we detected an excess of negative LD among pairs of low frequency NS variants compared to S variants (Fig 5, p<0.0005). We also observe an excess of negative LD among NS variants in 1KGP data when we match pairs on mean *B*-value amongst pairs of variants, and the chromosome each variant resides on.

Recent studies have identified batch effects present in the 1KGP that have led to rare population-specific artifacts [40]. We hypothesized that if the excess of negative LD amongst pairs of NS variants is due to batch effects, we would not detect this difference across multiple populations. However, we see qualitatively similar patterns in 4 of the 5 populations from the 1KGP. Additionally, previous work has suggested that low-frequency errors that are the consequence of batch effects tend to be in positive LD with each other [41]. Also, variants that are identified as error candidates are more likely to be NS variants [41]. With this in mind, we suspect that sequencing errors would bias our analysis of pairs of low frequency variants annotated as being NS towards being more often in positive LD than in negative LD. This suggests that our observation of a depletion of positive LD amongst NS variants is conservative. Lastly, the fact that we replicate these findings using unphased genotypes for H_R_^(2)^ and H_R_^(1)^ from high coverage sequencing suggests our conclusions are not due to artifacts in the 1KGP (Fig 6).

Our work adds to the growing literature indicating Hill-Robertson interference is a non-negligible force involved in the spatial distribution of NS variation in the human genome. Sohail *et al.* [18] found that putatively deleterious loss of function mutations were under-dispersed compared to putatively neutral S variation. Their summary statistic essentially quantified an increase in negative LD between loss of function variants. Their work is similar to our finding of an excess negative LD amongst NS variants compared to S variants. However, because the variants they examined were predominantly on different chromosomes and not physically linked, Sohail et al. mainly attributed this excess of negative LD as a signature of synergistic epistasis among deleterious variants. Hussin et al. [29] looked for signatures of interference by quantifying the relative enrichment of deleterious mutations in cold spots of recombination. Our simulations are concordant with this finding and predict that differences in LD amongst deleterious variants will be most distinguishable in regions of low recombination (*r*=1 x 10^-9^ per bp) and almost indistinguishable in regions of high recombination (*r*=1 x 10^-6^ per bp) (Fig 1).

Our results suggest that inference of demography that depends on genome-wide LD patterns may be biased if negative selection is not accurately modeled. For example, Tenesa et al. used *r*^2^ at different genetic distances to infer changes in population size and fit exponential functions to LD decay to infer admixture times [6]. If variants under negative selection are included in these analyses, they may bias parameter estimates. Further, demographic models fit to the site frequency spectrum in humans and *Drosophila* do not recapitulate empirically observed LD patterns [42–44]. Unmodeled negative selection may contribute to this lack of fit. We recommend carefully filtering variants that may be affected by selection. Alternatively, when considering variants under selection, we find that different combinations of recombination rate, DFE, and dominance predict unique mean r^2^ and *D* decay patterns. Thus, LD patterns may offer a strategy to infer the relationship between the dominance coefficient and the DFE in outcrossing populations [45]. However, in order to accurately predict the expected LD decay curve under a given scenario, extensive forward simulations are needed as a closed form solution for the predicted LD decay as a function of these genomic and selective parameters does not currently exist.

Further, through methods like LDscore regression, LD patterns have been used to learn about the architecture of complex traits [46]. We have shown that for a given allele frequency, variants under selection will have less LD as quantified by *r*^2^ than putatively neutral variants. This effect may bias methods that assume variants of the same allele frequency will have the same distribution of *r*^2^ values. For example, stratified LD score regression is used to infer heritability in different functional annotations and assumes that there will be similar LD between causal variants and tag SNPs regardless of the fitness effects of the causal variants [47]. Because causal variants under greater negative selection may have larger effects on the trait [48], the heritability that deleterious variants account for may be systematically underestimated.

Epistatic interactions have been documented between variants in different genes [49] as well as variants within the same gene [50] and are believed to be widespread properties of biological networks [51, 52]. Although the genomic consequences of additive deleterious variants which multiplicatively (across loci) affect fitness are extensively studied, pairwise interactions amongst variants might be a non-negligible force governing molecular evolution. Current methods to identify plausible pairwise interactions between SNPs rely on classic population genetic summary statistics of linkage disequilibrium or other summaries of pairwise association frequencies [51,53,54]. Although we detect differences in LD summary statistics in the human genome amongst variants of different annotations, our simulations suggest this difference can be explained by interference without the need to invoke epistasis (Fig 7). We hypothesize that the difference is predominantly due to interference because of its relationship with recombination. Our simulations and theory predict that the biggest differences in LD should be present amongst variants with the least amount of recombination separating them, and the smallest differences in LD should be present amongst variants with the most amount of recombination separating them. This scenario aligns with what we observe in our empirical data set. We propose that when developing methods for detecting synergistic epistasis, null models should incorporate Hill-Robertson interference.

Future work could quantify the prevalence of epistasis amongst linked deleterious variants while jointly quantifying the fitness effects of pairs of deleterious variants. Additionally, although it has been shown that two locus statistics such as *D*’ can be used in the detection of epistatic interactions and interference [19, 55], it is possible that a combination of summary statistics can better discern between the two. Machine learning approaches, which can combine different features of genetic variation data [56], may provide a powerful tool for detecting salient spatial features of proximal variants involved in epistatic interactions and interference.

## Materials and Methods

### Forward Simulations

The distribution of genomic elements in our forward simulations followed the specification in the SLiM 4.2.2 manual (7.3), which is modeled after the distribution of intron and exon lengths in Deutsch and Long [57]. Within exonic regions, NS and S mutations were set to occur at a ratio of 2.31:1 [58]. For simulations using a DFE, the selection coefficients (*s*) of NS mutations were drawn from a gamma-distributed DFE with shape parameter 0.186 and expected selection coefficient E[s] = −0.01314833 [59]. All NS mutations were either additive with *h*=0.5 or recessive with *h*=0.0. The per base pair per generation recombination rate was constant across each simulation region and was fixed at either r ∈{10^−6^, 10^−7^, 10^−8^, 10^−9^} while the per base pair per generation mutation rate was set to µ=1.5 x 10^−8^. No simulation parameters were scaled.

### Empirical Data

We used data from 50 random Yoruban (YRI) individuals from the 1KGP. Specifically, we randomly sampled 50 individuals from Supplementary Table 4 in Gazal *et. al* [60] that were labeled as coming from the “YRI” population, had a mating type described as “OUT” (outcrossing), and had a Q-score (quality score) of greater than 50. Then we removed all non-biallelic variants. Next, we polarized the remaining variants using only high confidence sites in the 6-way primate EPO multiple alignment as the ancestral allele. All variants without a high-confidence ancestral allele were also removed from analysis. Remaining biallelic exonic variants in our sample were annotated as either NS or S variants using ANNOVAR [61]. For analysis with different populations of the 1KGP Phase 3 data, the same procedure described above was used.

Analysis with the YRI NYGC 1KGP resequenced data used the same 50 individuals that were selected from the 1KGP Phase 3 data. Variants were also polarized using only high-confidence sites in the EPO multiple alignment as the ancestral allele. All variants without a high-confidence ancestral allele were also removed from analysis. Remaining biallelic exonic variants in our sample were annotated as either NS or S variants using ANNOVAR [61]. Pairs of variants that were within 10,000 bp of each other, had the same allele count in the sample of 50, had the same annotation, and were both located in the 1KGP strict mask (http://ftp.1000genomes.ebi.ac.uk/vol1/ftp/data_collections/1000_genomes_project/working/20160622_genome_mask_GRCh38/StrictMask/20160622.allChr.mask.bed) were then analyzed.

### Computing LD summary statistics with frequency matching

Three different pairwise LD statistics were calculated between SNPs with the same allele count in our sample. First, to compute *D* we used the formula from Lewontin (1964):

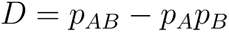

Here, *D* is the difference between the observed frequency of haplotype *AB* (*p_AB_*) and the expected frequency of haplotype *AB* (assuming random association of the alleles at the two loci, *p_A_*p_B_*). In this calculation, *p*_A_ and *p_B_* are the observed frequencies of the derived alleles *A* and *B* in our sample. With the notation described here, the coupling haplotypes include derived alleles at both variants (*AB*) and the repulsion haplotypes include derived alleles at one variant (*Ab* are *aB*).

We computed *r*^2^ with the formula:

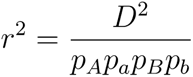

*D*’ was computed with:

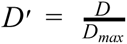

where:

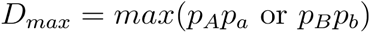

Whether *D*’ is negative or positive depends on the arbitrary choice of the alleles paired at two loci. We chose the pair of derived alleles to be the pair of alleles that cause *D*’ > 0 when located in coupling. Therefore, a pair of derived doubletons that only ever appear in a sample together on the same haplotype will have a *D*’ = 1. A pair of derived doubletons that are never observed on a haplotype together in a sample will have a *D*’ = −1 (**SI Fig 3**).

Limiting LD calculations between SNPs to restricted allele frequency intervals was first done by Eberle et al. [31] and found to be a more sensitive measure for assessing the average decay of LD and is able to generate average *r*^2^ values across nearly the entire informative range. We applied this approach to pairs of NS SNPs as well as pairs of S SNPs. After LD statistics were computed for NS variants, the same method described above was followed for S variants. The decay of these LD summary statistics were then modeled between pairs of variants with a smooth line generated using a generalized additive model using the “gam” function from the R package *mgcv* version 1.8-29 [62]. We fit the default formula y ∼ s(x, bs = “cs”).

### A new LD statistic for unphased genotypes: H_R_^(j)^

To summarize the counts of homozygous ancestral, singly heterozygous, and doubly heterozygous genotypes amongst pairs of NS and S variants we took two approaches. The first approach was taken with both simulated data generated with SLiM and 1KGP Phase 3 data. Genomic data from these sources are already phased. We first used custom scripts to unphase the data sets, then we computed the distribution of counts of homozygous ancestral, singly heterozygous, and doubly heterozygous genotypes amongst pairs of variants that were within 10,000bp from each other, had an identical allele count, and also had the same functional annotation (S or NS). Here variants can have three genotypes 0 (homozygous ancestral), 1 (heterozygous), 2 (homozygous derived). For example, for a pair of NS singletons in a sample of 50 diploid individuals, the counts could be: n_11_ = 1, n_01_ and n_10_ =0, and n_00_=49. This would correspond to 1 individual being doubly heterozygous for this pair of variants, 0 individuals being singly heterozygous, and 49 individuals being homozygous for the ancestral genotypes at these loci. The second approach is identical to the first, however, for the YRI NYGC 1KGP resequenced data, the data was not phased to start with. Therefore, no custom scripts were used to unphase the data.

We developed a new statistic to be applied to unphased genotype data to quantify whether derived deleterious alleles at a pair of SNPs are likely to be found on the same haplotype. We call our new static H_R_^(j)^:

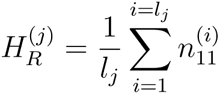

H_R_^(j)^ depends on three components: n^(i)^_11_, j, and l_j_. *j* refers to the allele count of variants to be analyzed. For singletons, *j* = 1, for doubletons *j* = 2, for tripletons *j* would be equal to 3, etc. *l*_j_ is the number of pairs of variants at allele count *j* within a distance threshold. *n*^(i)^_11_ represents the count of heterozygous individuals with derived variants at both loci of the pairwise comparison *i* (coding genotypes as 0 for homozygous ancestral, 1 for heterozygous, 2 for homozygous derived). For pairs of doubletons, this equation is represented as:

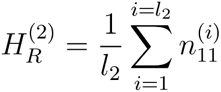

To compute H_R_^(2)^ for derived NS variants in a sample we created a list of all derived NS doubletons in the sample. Second, we created a list of all unique pairwise combinations of these doubletons that involve SNPs within a certain distance threshold. The length of this list would be equal to *l*_2_. We then find the number of heterozygous individuals with derived variants at both loci of the pairwise comparison *i* are then summed together over all *l*_2_ pairs of SNPs. This is represented by 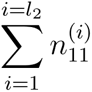. Lastly, division by the total number of pairwise comparisons (or multiplication by 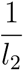) gives H_R_^(2)^.

In principle, H_R_^(j)^ can be computed for any value of *j*<2*n*, where *n* is the number of individuals in the sample. In practice, we consider *j*=1 and *j*=2 because low-frequency variants are in general most likely to be impacted by negative selection (**SI Fig 2**).

### Estimating Genetic Distance Between Variants

The genetic distance between two markers was computed using the high-resolution pedigree-based genetic map assembled by deCODE [63]. First, we averaged the male and female genetic maps. Occasionally the genetic distance between two markers in our sample was not explicitly estimated by deCODE. If one or both of the markers were in regions not measured by deCODE we removed these markers from our analysis with genetic distance. However, if the two markers were within a region of the genome with a high-resolution recombination environment estimated by deCODE, we imputed the genetic distance between markers by estimating a centiMorgan per base pair rate and multiplying this rate by the physical distance between the two markers that lacked a genetic distance annotation.

### Matched-pairs permutation test for differences in LD between S and NS SNPs

To quantify differences in LD while controlling for possible covariates, we conducted a matched-pairs permutation test on pairs of SNPs. Using the R Package “MatchIt” [64], for each pair of NS SNPs, we extracted a pair of S variants with a similar physical distance between variants, the same allele count amongst pairs, a similar genetic distance (cM), similar mean *B*-value amongst pairs of variants, and located on the same chromosome as a NS pair of variants in our sample. **S3 Fig** shows how closely we required the pair of S SNPs to match the pair of NS SNPs across these characteristics. We then computed the difference between the LD statistic for the NS pair and the S pair. The mean of these differences across all the matched pairs was then computed and used as the summary statistic (dashed line in Figure 5B). A matched-pairs permutation test was then conducted with 10,000 permutations. In each test, the null hypothesis was that the mean difference in LD statistics between pairs of NS and S SNPs was 0. The alternative hypothesis was that the mean difference in LD statistic across pairs was less than 0 (the LD statistic among NS pairs was less than the matched S pair). A graphical representation of this method can be seen at **SI Fig 10.**

### Annotating variants with amount of background selection

For one of our matched-pairs permutation tests, we matched pairs of NS variants with pairs of S variants with similar levels of background selection. *B*-values were downloaded from http://www.phrap.org/software_dir/mcvicker_dir/bkgd.tar.gz. Then we used liftOver (UCSC Genome Browser) with its default settings and the hg18Tohg19 chain (http://hgdownload.cse.ucsc.edu/goldenPath/hg18/liftOver/hg18ToHg19.over.chain.gz) to convert *B*-value coordinates from hg18 to hg19. Each SNP in our data set was then annotated with its corresponding *B*-value using the GenomicRanges package [65].

## Acknowledgements

This work was supported by NIH grant R35GM119856 to KEL and a Gates Foundation Fellowship to JAG. We thank Nandita Garud and Jazlyn Mooney as well as members of the Lohmueller lab for helpful discussions throughout the project and comments on the manuscript. The 30x coverage sequencing of the 1000 Genomes samples was generated at the New York Genome Center with funds provided by NHGRI grant 3UM1HG008901-03S1. We would like to thank the New York Genome Center for producing and making these data available pre-publication.

