## Supplementary Information for "Negative linkage disequilibrium between amino acid changing variants reveals interference among deleterious mutations in the human genome"

#### Supporting Information

##### SI Fig 1

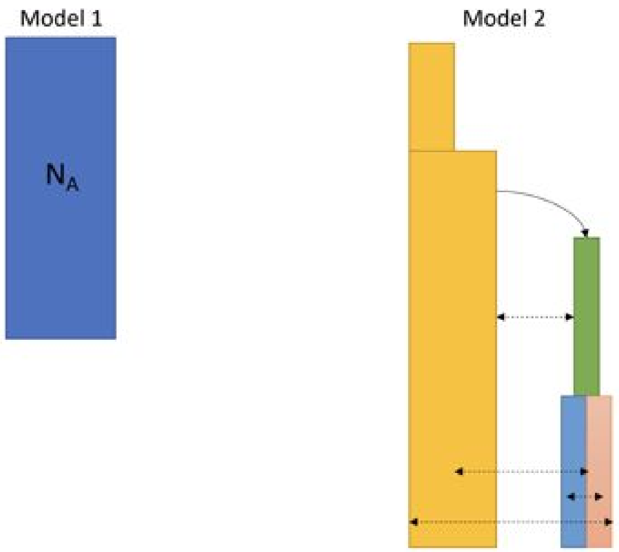


**Fig S1. Demographic models for simulations.** Model 1 represents a constant population size simulation. Model 2 represent the Gravel et. al (2011) demographic model. The yellow subpopulation represents Africans (YRI). The green represents the ancestral Eurasian bottleneck. The blue represents East Asians and the pink represents Europeans. Model 3 is the Gravel demographic model but without migration.

**SI Fig 2**

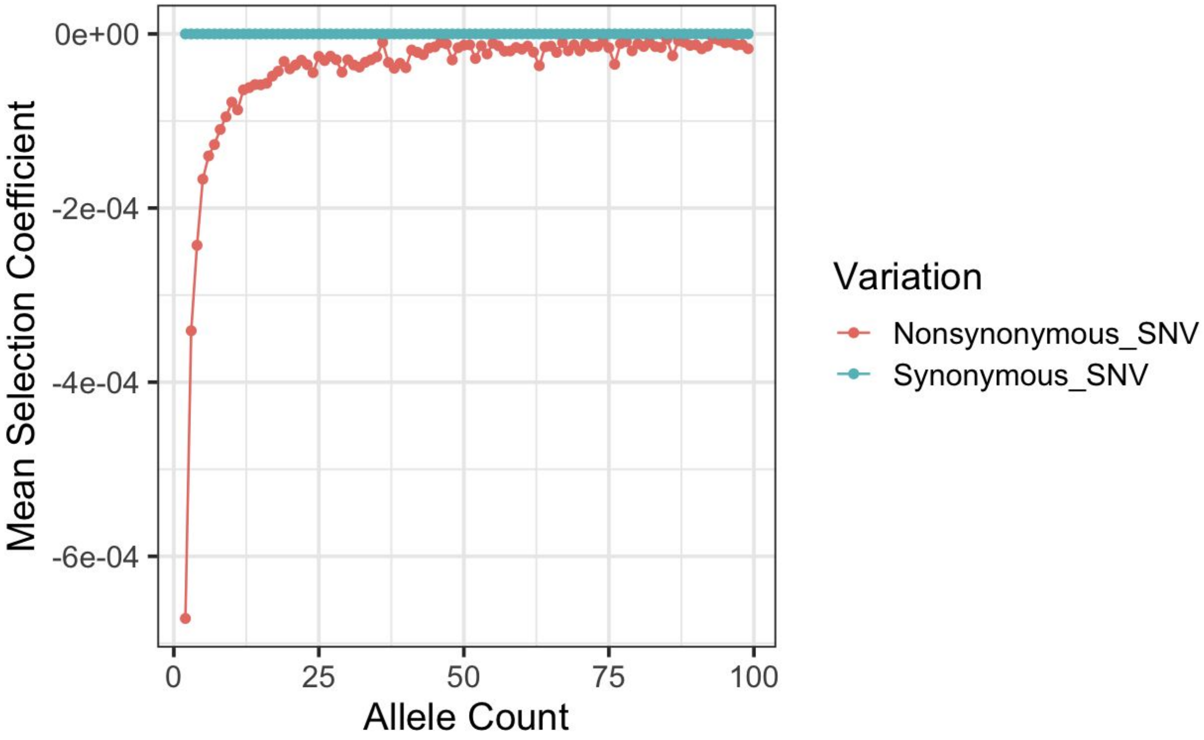


**Fig S2. Doubletons in a sample of 50 individuals are more deleterious than higher frequency variants.** We simulated 300 replicates of a constant population size of 14, 474 diploid individuals with $r=1 \times 10^{-8}$ per bp, and the DFE and genome structure defined in Methods. Although simulations predict singletons (first red point) on average should be the most deleterious in our samples, doubletons (second red point) are also relatively deleterious. Higher frequency variants tend to have mean selection coefficients that are much more neutral. Because we want to study the effects of negative selection on LD, we restrict many of our analyses to low frequency variants (allele count <= 5)

**SI Fig 3**

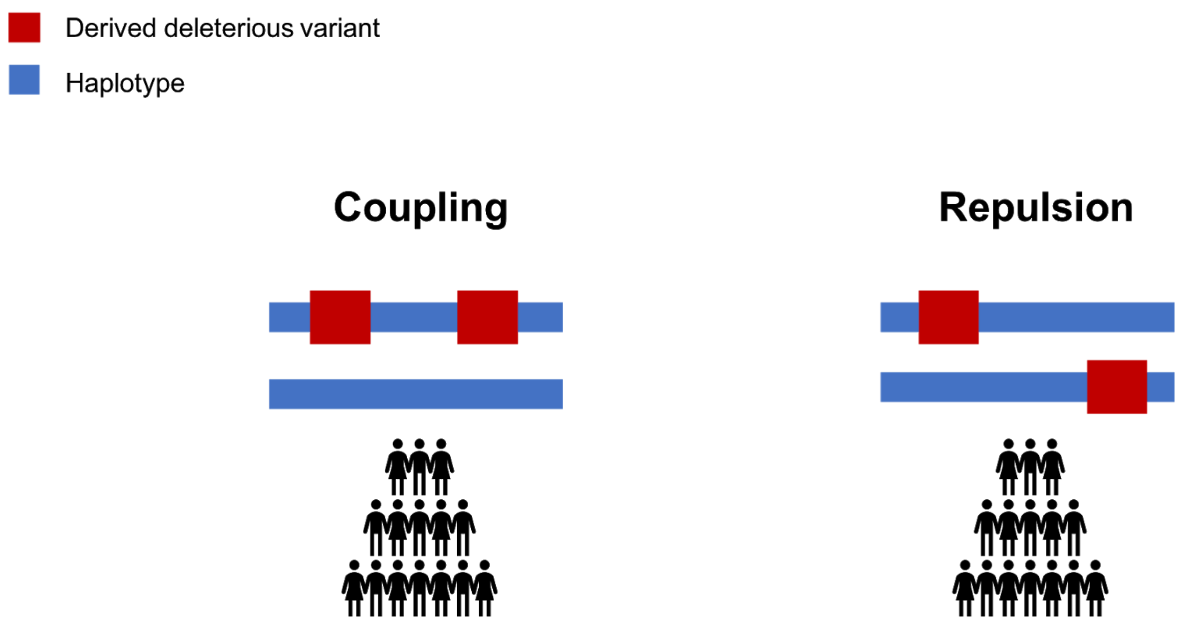


**Fig S3. Pictorial representation of variants in coupling and repulsion.** In this study, pairs of derived variants that co-occur on the same haplotypes more frequently than expected are said to be "coupling" or in positive LD (left panel). Pairs of derived variants that occur less frequently than expected on the same haplotype are said to be in "repulsion" or in negative LD (right panel).

**SI Fig 4**

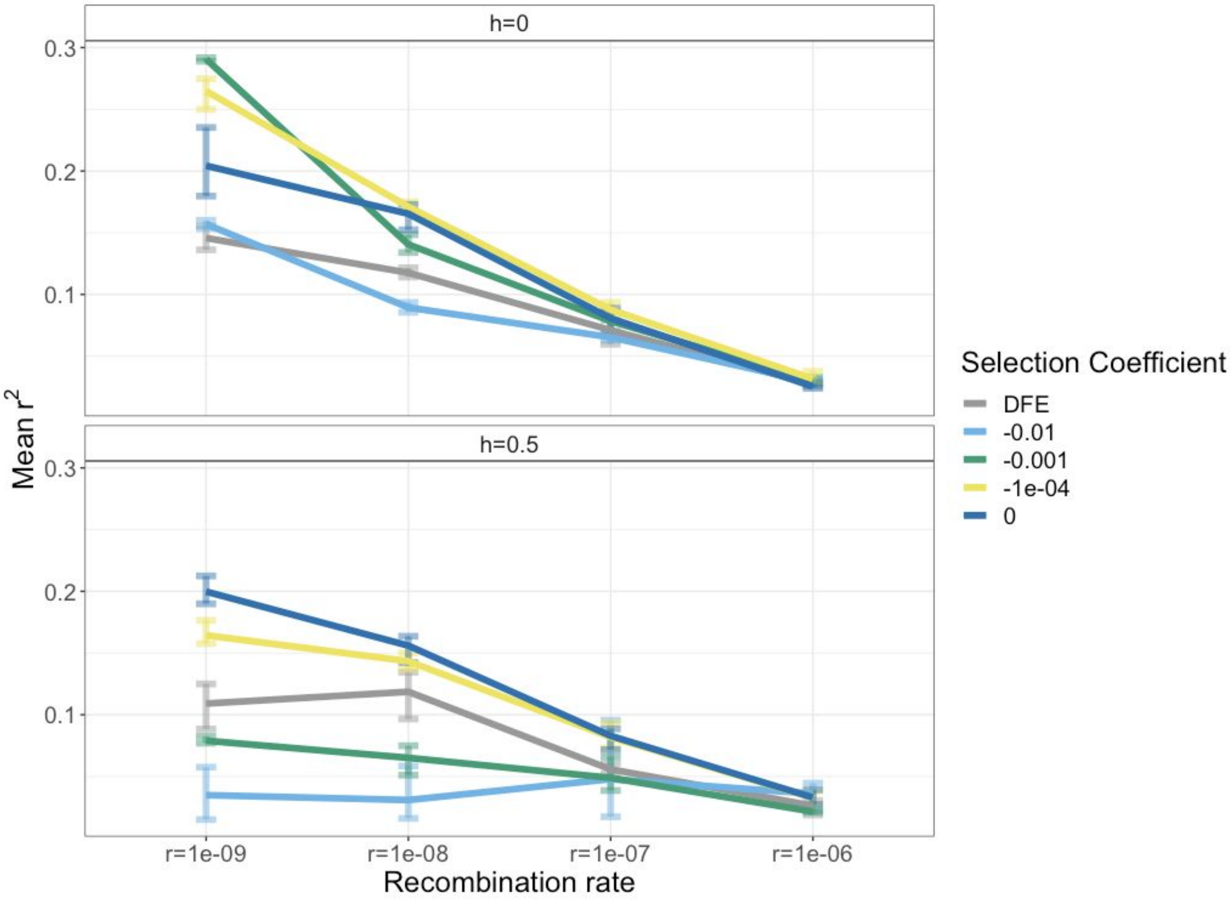


**Fig S4. Mean $r^2$ for simulated nonsynonymous doubletons 10kb from each other across different recombination and dominance parameters.** The differences in the decay curves are most apparent for recombination rate r=1 x $10^{-9}$ per bp and depend on the dominance coefficient (*h*) and the selection coefficient of NS mutations.

**SI Fig 5**

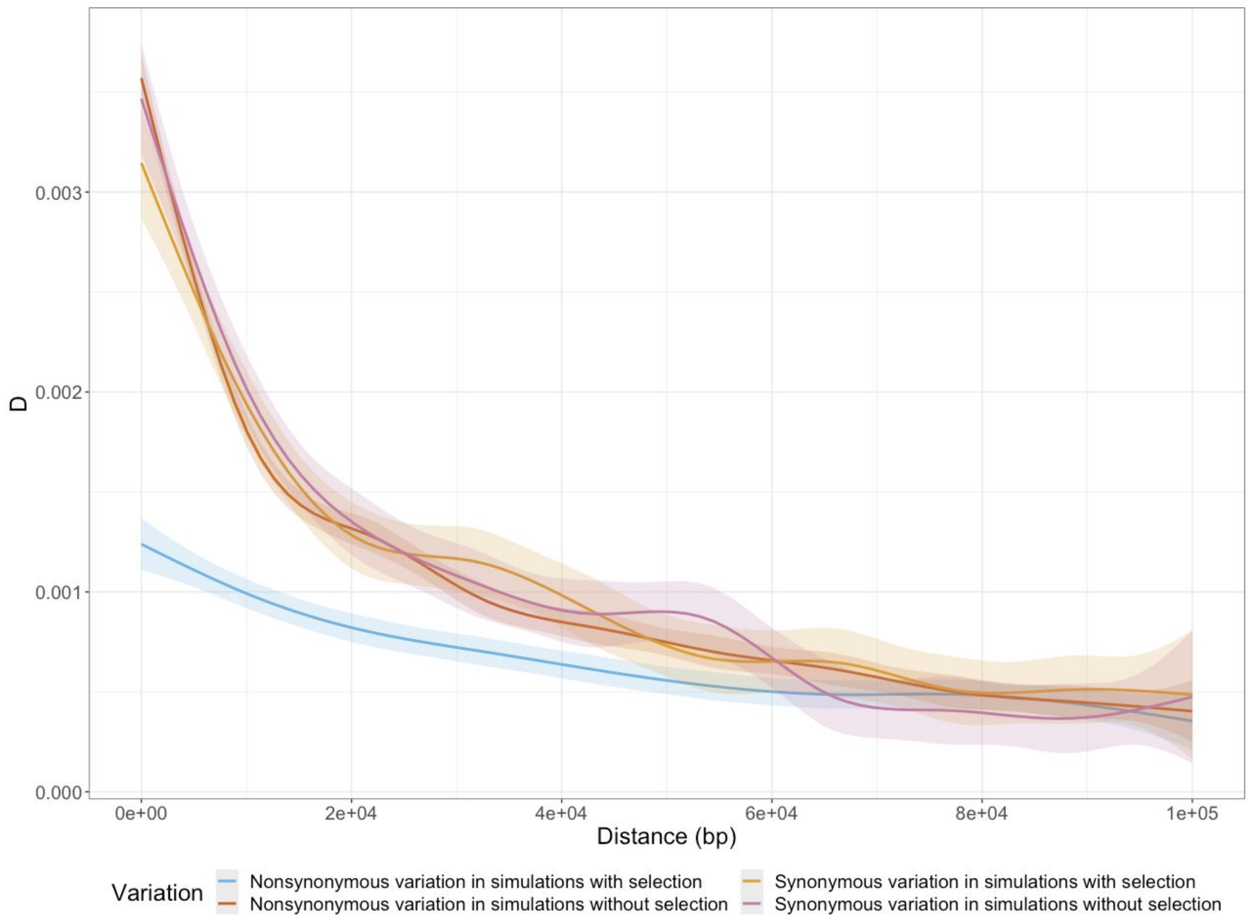


**Fig S5. Decay of *D* from simulated NS and S doubletons.** There are two different scenarios visualized in the figure: 1) simulations with negative selection, and 2) simulations without negative selection (completely neutral evolution). The blue line is the decay curve of NS doubletons with direct negative selection acting on them. In contrast to the decay curve of the NS doubletons not under negative selection (dark red), the NS doubletons with direct negative selection acting on them (blue) have a more negative *D* than the other neutral doubletons (S variants in simulations without selection, NS variants in simulations without selection, S variants in simulations with selection).

**SI Fig 6**

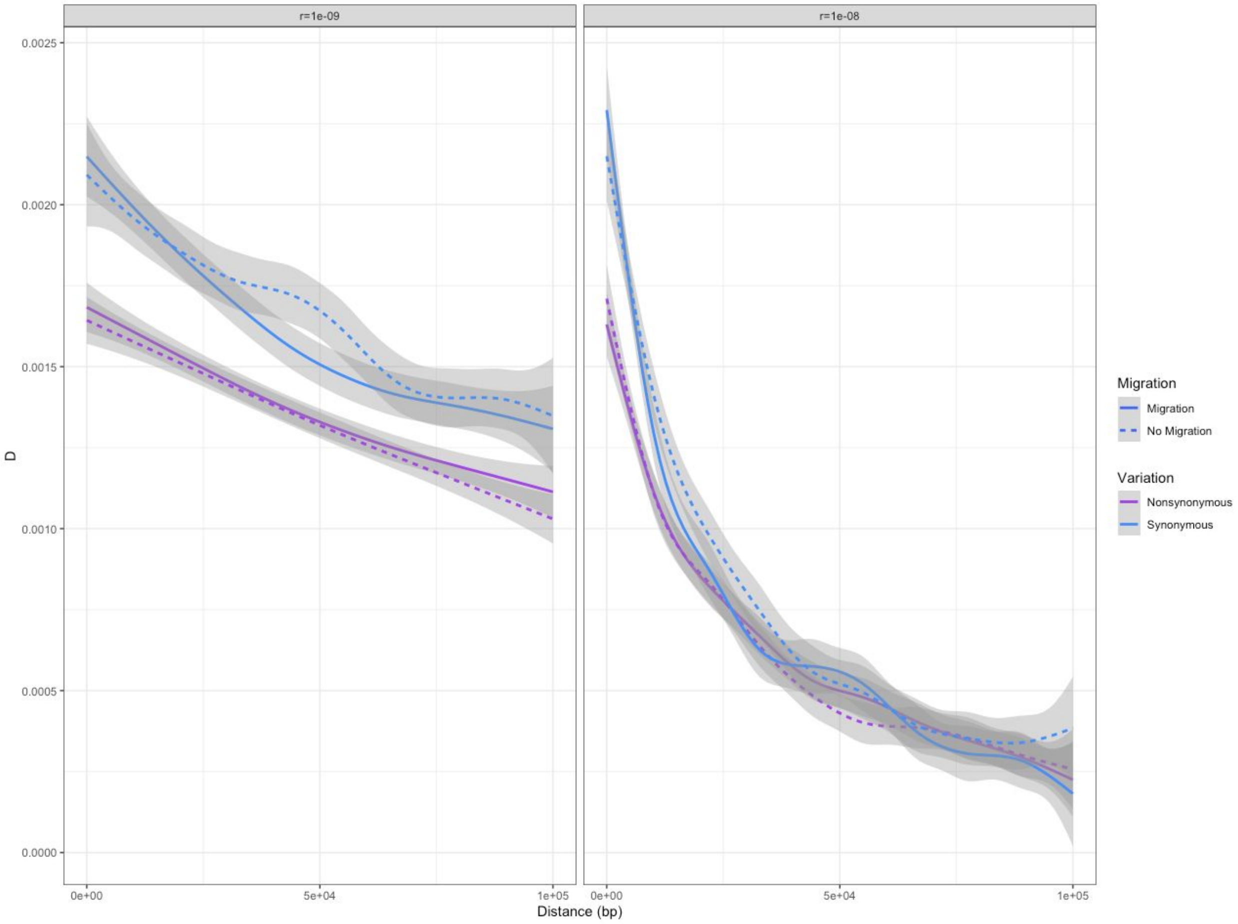


**Fig S6. Decay of *D* for simulated NS and S doubletons in scenarios with migration.** For *r*=1 x $10^{-8}$ per bp, the largest differences between S and NS *D* is predicted by our simulations to be within the 0-10,000 bp range. In our simulations with *r*=1 x $10^{-9}$ per bp, noticeable differences between S and NS *D* is predicted by our simulations to be across the 0-100,000 bp range. Additionally, migration appears to qualitatively obscure the difference in *D* between types of variants at intermediate distances.

**SI Fig 7**

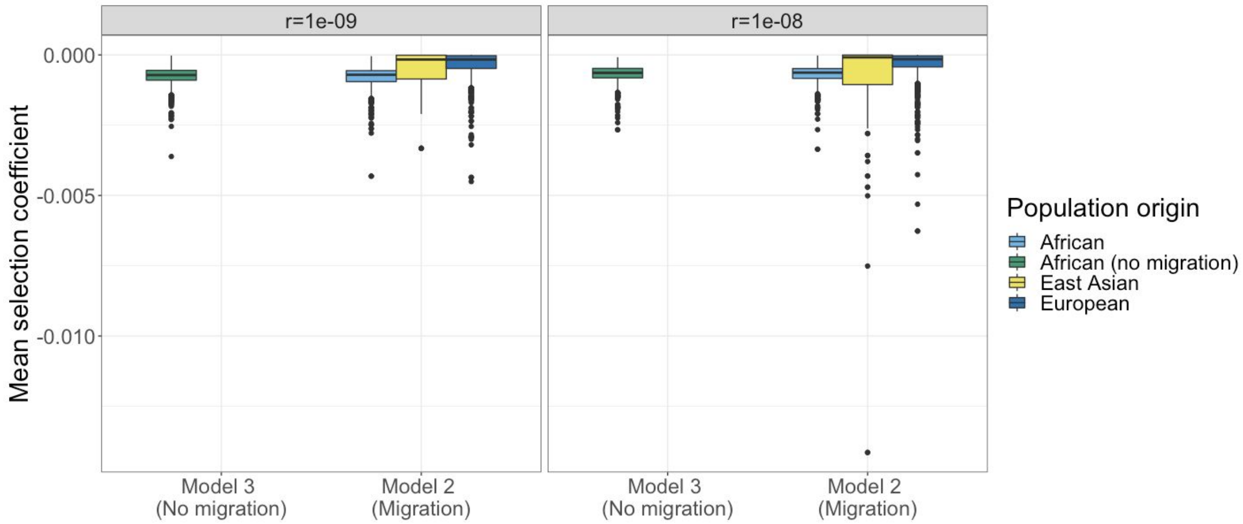


**Fig S7. Mean selection coefficient of deleterious doubletons in our simulated samples and their population of origin.** For both $r$=1 x $10^{-9}$ per bp and $r$=1 x $10^{-8}$ per bp, doubletons that appear in our simulated "African" sample, have on average different mean selection coefficients depending on their population of origin. Migration (Model 2) allows for deleterious variants that originated from other populations to appear in our African samples. On average, the doubletons in our sample of African individuals that originate from East Asia and Europe are less deleterious than the doubletons in our sample that originated from Africa.

**SI Fig 8**

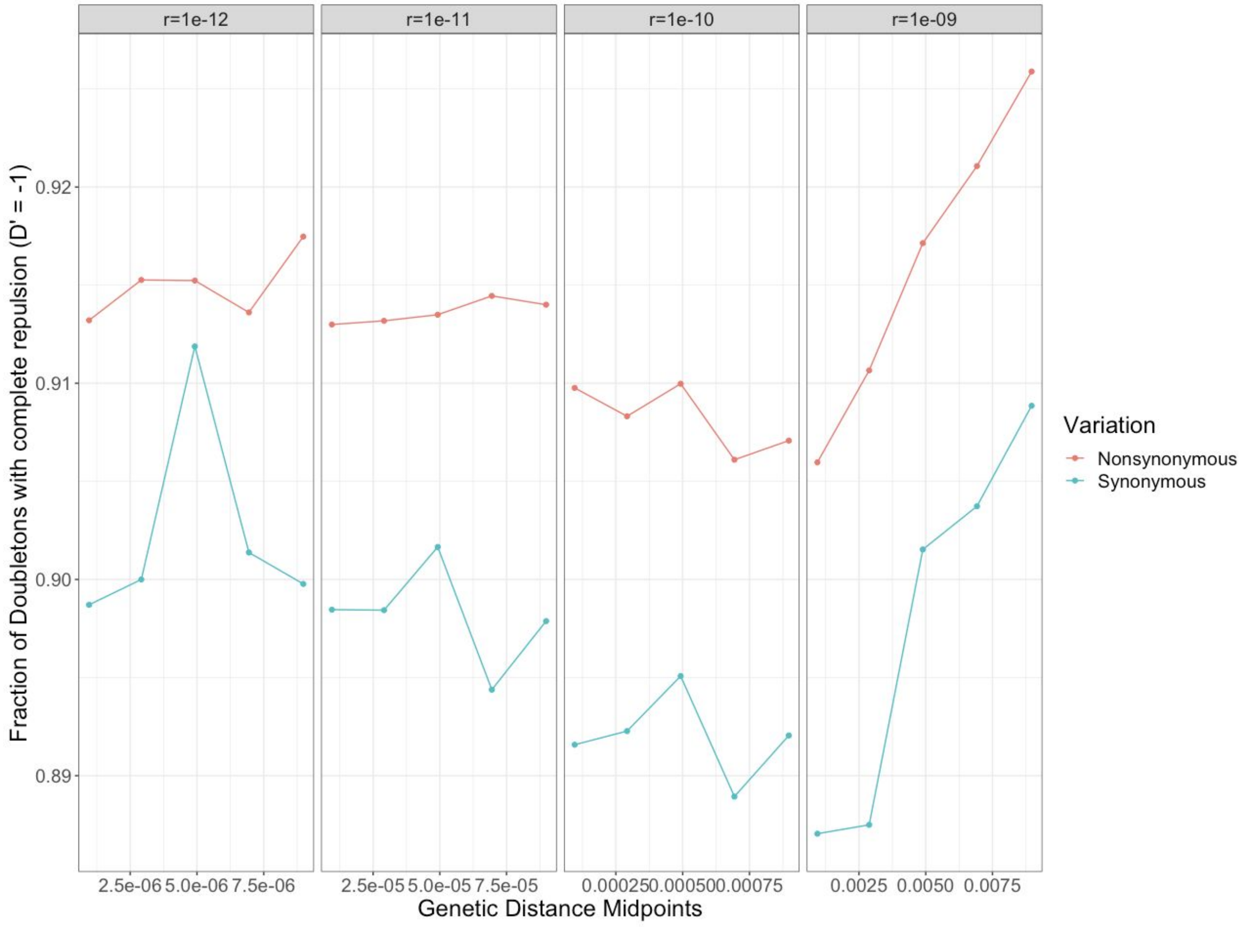


**Fig S8. Simulations predict NS pairs of doubletons should be more often in complete repulsion than S pairs of doubletons.** The red line denotes the total fraction of simulated NS doubletons that are in complete repulsion (*D'* = -1) and the blue line denotes the total fraction of simulated S doubletons that are in complete repulsion.

**SI Fig 9**

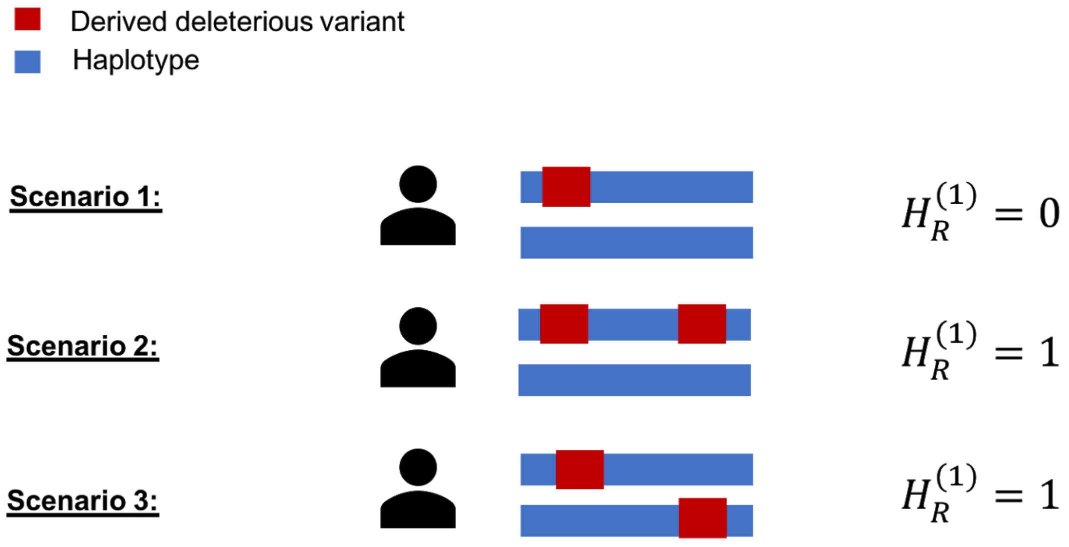


**Fig S9. Pictorial representation of $n_{AB}$ and** $H_R^{(1)}$**.** The statistic $H_R^{(1)}$ depends on the number of unique pairwise comparisons amongst doubletons within the distance threshold 10kbp ($l_1$=1). Additionally, it depends on the number of individuals who are heterozygous at both loci. In both Scenario 2 and 3, the individual is heterozygous at both loci, thus making $H_R^{(1)}$= 1. The statistic $n_{AB}$depends on the number of haplotypes that contain derived alleles at loci. In this case, only Scenario 2 contains one haplotype that has both derived variants. $H_R^{(j)}$ can also be thought of as an indicator variable taking the value of 1 if an individual is heterozygous at both loci and 0 if not.

**SI Fig 10**

<u>Complete List of Pairwise LD Computations</u>

| Allele Count | Position of SNP 1 | Position of SNP 2 | Distance | Chromosome | D | Annotation |
|---|---|---|---|---|---|---|
| 3 | 148832 | 149832 | 1,000 | 2 | 1 | Synonymous |
| 4 | 700005 | 710005 | 10,000 | 3 | .2 | Nonsynonymous |
| 3 | 452380 | 452395 | 15 | 5 | 0.5 | Nonsynonymous |
| 1 | 169832 | 178368 | 8,536 | 2 | 1 | Synonymous |
| 3 | 745002 | 746004 | 1,000 | 2 | 0 | Nonsynonymous |
| 4 | 300000 | 310000 | 10,000 | 3 | 0.5 | Synonymous |
| 3 | 503000 | 506000 | 3,000 | 5 | 0 | Nonsynonymous |
| 1 | 149832 | 158368 | 8,536 | 2 | 1 | Synonymous |

<u>Matched SNPs</u>

| Allele Count | Position of SNP 1 | Position of SNP 2 | Distance | Chromosome | D | Annotation |
|---|---|---|---|---|---|---|
| 3 | 745002 | 746004 | 1,000 | 2 | 0 | Nonsynonymous |
| 3 | 148832 | 149832 | 1,000 | 2 | 1 | Synonymous |
| 4 | 700005 | 710005 | 10,000 | 3 | 0.2 | Nonsynonymous |
| 4 | 300000 | 310000 | 10,000 | 3 | 0.5 | Synonymous |

Observed Mean Difference in D Between Matched Pairs:

$$\frac{(0\text{-}1) + (0.2\text{-}0.5)}{2} = -0.65$$

**Fig S10. Flow chart of matched-pairs permutation test.** First, sample 50 individuals from a population. Polarize variants and then annotate variants as either NS or S. Second, compute LD summary statistics among pairs of variants that are within 10,000 bp from each other and have the same allele count (AC) and annotation (NS or S). Third, for each pair of NS variants with a computed LD statistic, find one S pair of variants with the same AC, on the same chromosome, and with a similar (<50 bp) distance between variants. Each pair of NS and S pairs constitutes a matched pair. Fourth, compute the mean difference between matched pairs. Fifth, permutate the label (i.e. S or NS) for each pair of SNPs

**SI Fig 11**

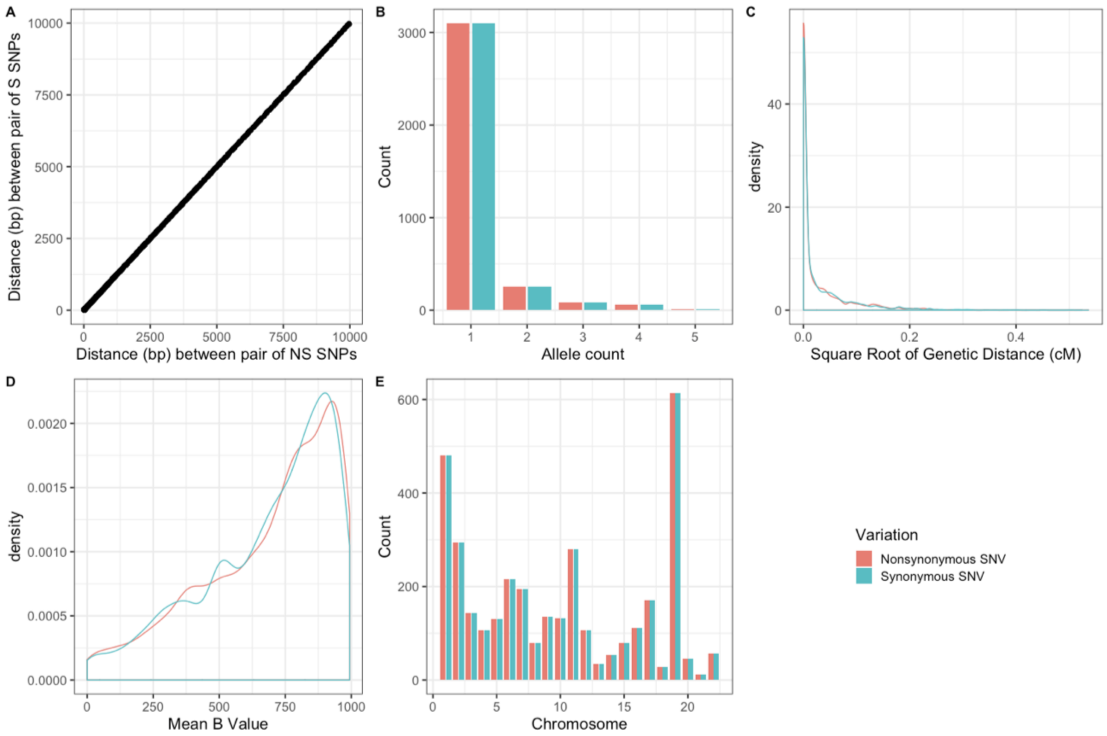


**F**

| Feature | Matching | NS Mean | NS SD | S Mean | S SD |
|---|---|---|---|---|---|
| Distance (bp) | Within 50 bp | 2344.52 | 2852.937 | 2344.607 | 2852.929 |
| Allele count | Exact matching | 1.181039 | 0.5778994 | 1.181039 | 0.5778994 |
| Mean B value | Within 200 units | 684.8885 | 251.2354 | 681.1461 | 246.2606 |
| Chromosome | Exact matching | 9.817818 | 6.742537 | 9.817818 | 6.742537 |
| Genetic distance | NA | 0.003472501 | 0.01318808 | 0.0035600 | 0.01305732 |

**Fig S11. Matching rules for features and their distribution across NS variation and S variation in empirical data.** In order to reduce possible confounding effects of different distances, allele counts, levels of background selection, and recombination rate, we implemented a matched pairs scheme. For each pair of NS variants in our data set, we attempted to identify a pair of S variants with similar criteria. This allowed us to compare pairs of variants with similar distributions and spread. The "Matching" column denotes the rules we used for matching. "SD" stands for standard deviation.

**SI Fig 12**

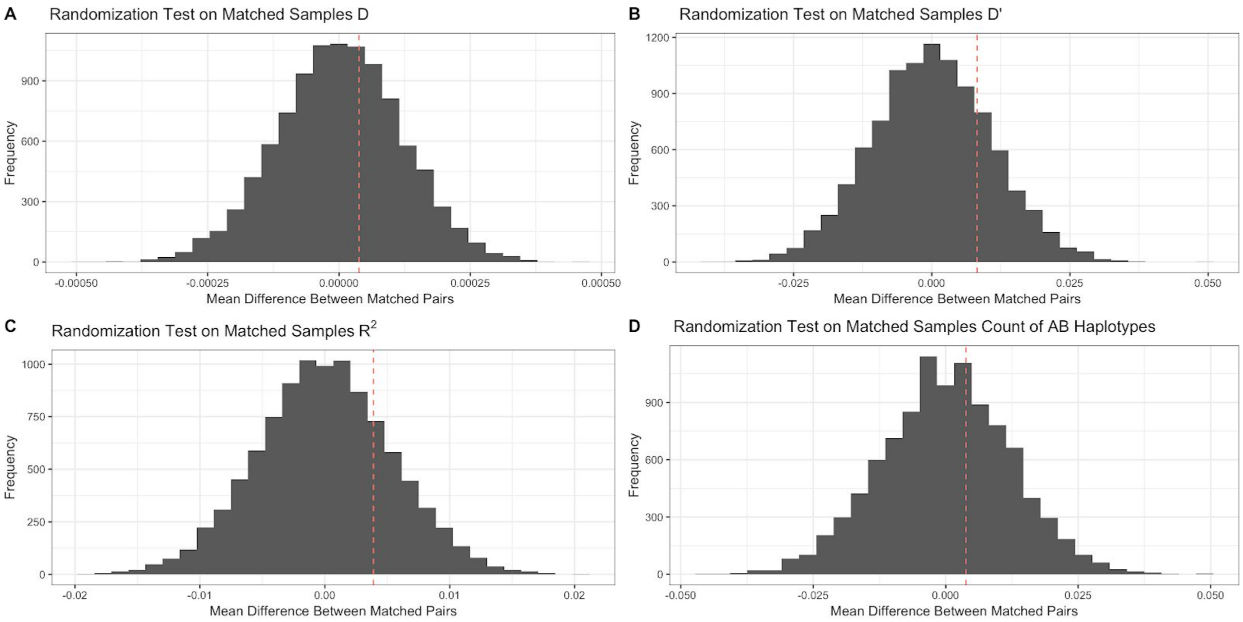


**Fig S12. Matched-pairs permutation test on various LD summary statistics in simulations without negative selection**. In simulations without negative selection, none of the LD summary statistics we computed showed a significant difference between NS and S matched pairs.

**SI Fig 13**

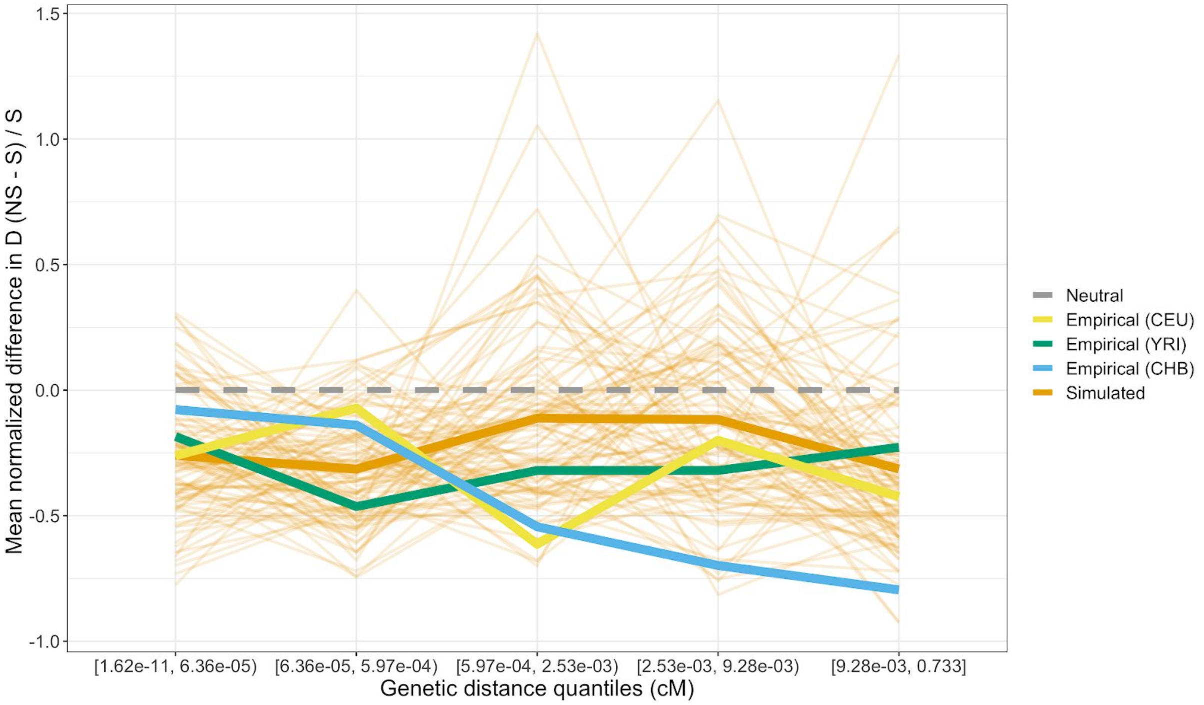


**Fig S13. Mean normalized difference in *D* across genetic distance quantiles for non-African 1KGP populations.** Empirical (green) and simulated data shows a deficit in *D* between NS variants. The lighter orange lines show 100 resamples of the simulated data. Each resample of the simulated data has the same amount of variants with the same allele count as the YRI empirical data. Both CHB and CEU also show a more negative *D* between low frequency NS variants compared to low frequency S variants.

**SI Fig 14**

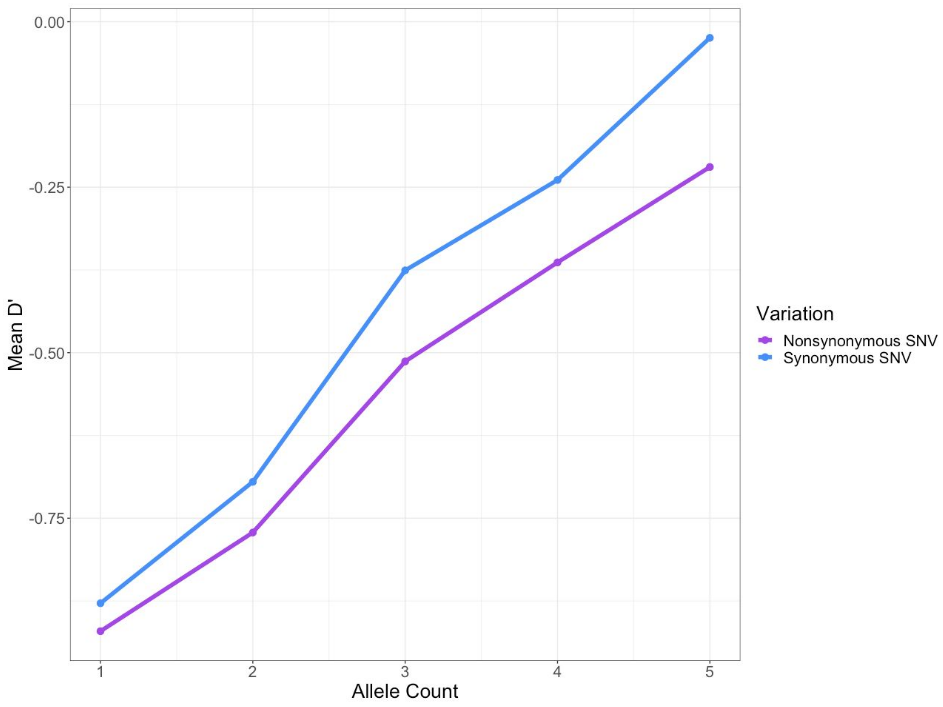


**Fig S14. Mean *D*’ of matched pairs of simulated NS and S SNPs across allele counts in a sample of n=50 diploid individuals.** On average, more negative *D*’ is seen across pairs of low frequency NS polymorphisms. In this analysis, the recombination rate was set to a constant 1 x $10^{-8}$ per bp with a constant population size of 14,474 diploid individuals. 300 simulation replicates were then aggregated and the mean *D*’ was computed for each frequency.

### SI Tables

**SI Table 1**

Summary Statistics of D' Across Allele Counts (Simulated Data With Selection)

| | | Nonsynonymous Pairs | | Synonymous Pairs | |
|---|---|---|---|---|---|
| Allele Count | Number of Matched Pairs | Mean | SD | Mean | SD |
| 1 | 7490 | -0.921 | 0.390 | -0.879 | 0.478 |
| 2 | 810 | -0.772 | 0.635 | -0.695 | 0.718 |
| 3 | 189 | -0.513 | 0.861 | -0.376 | 0.929 |
| 4 | 88 | -0.364 | 0.937 | -0.239 | 0.968 |
| 5 | 41 | -0.220 | 0.988 | -0.024 | 1.012 |

**S1 Table. Summary statistics of *D'* across allele counts for neutral (S) and deleterious (NS) variants.** In simulations, NS pairs of doubletons are predicted to have a lower *D'* compared to S pairs of doubletons.
